# Intermittent fasting mitigates cognitive deficits in Alzheimer’s disease via the gut-brain axis

**DOI:** 10.1101/2022.05.11.491466

**Authors:** Mengzhen Jia, Lin Shi, Yihang Zhao, Xinyu Hu, Junhe Zhao, Chen Ding, Yiqiu Shang, Xin Li, Xin Jin, Xiaoshuang Dai, Xuebo Liu, Zhigang Liu

## Abstract

Intermittent fasting (IF) is suggested to mitigate cognitive impairment in Alzheimer’s disease (AD). We detailed the effects of IF in AD development and assessed the contribution of the gut microbiota-metabolite-brain axis in promoting the effects of fasting. A 16-week IF regimen significantly improved spatial memory and cognitive function in AD transgenic mice; this improvement was accompanied by decreased Aβ accumulation and suppression of neuroinflammation. IF reshaped the microbiota and beneficially altered the microbial metabolites related to cognitive function. Multi-OMICs integration demonstrated a strong connection between IF induced changes in hippocampus gene expression, the composition of the gut microbes, and the serum metabolites beneficial for cognitive function. Removal of gut microbes with antibiotics abolished the neuroprotective effects of IF. These findings suggest new avenues for therapy and nutritional intervention to prevent the development of AD and other neurodegenerative diseases.

## Introduction

Alzheimer’s disease (AD) is a common neurodegenerative disease and the major cause of dementia, which affects 43 million people worldwide (Arbo et al., 2019). AD is characterized by cognitive impairment resulting from the accumulation of amyloid-beta peptides (Aβ) and intracellular neurofibrillary tangles caused by hyperphosphorylated Tau protein in the brain (Arbo *et al*., 2019; Eimer and Vassar, 2013). In addition, AD leads to synaptic damage and neuroinflammation in the brain (Kim et al., 2019). However, the etiological mechanism of AD is largely unknown. Medical treatments and effective interventions to prevent the development of AD are extremely lacking and urgently needed. The risk of AD is associated with genetic factors, obesity, lack of physical exercise, and an unhealthy diet (Silva et al., 2019).

Beneficial effects of dietary restriction on brain health and dementia are reported widely (Wahl et al., 2016). Similarly, beneficial effects of IF were reported for metabolic syndrome and cognitive impairment (Cherif et al., 2016). Specifically, by inducing biochemical and metabolic changes, intermittent fasting (IF) improved the cognitive function of elderly individuals at 36 months follow-up (Ooi et al., 2020). Animal studies suggested that IF could alleviate memory deficits by increasing the expression of neurotrophic factors, improving synaptic plasticity, and inhibiting neuroinflammatory responses (Hu et al., 2017; Liu et al., 2019; Liu et al., 2020; Vasconcelos et al., 2014). A recent study showed that a three-month IF regimen enhanced spatial cognition and hippocampal neurogenesis in female mice (Dias et al., 2021). Notably, IF reduced Aβ accumulation, a key indicator of AD development, and prevent memory loss in ovariectomized rats infused with β-amyloid (Shin et al., 2018; Zhang et al., 2017). In addition, in transgenic AD mouse models, IF conferred increased hippocampal neuron resistance to the toxic effects of kainic acid (Bruce-Keller et al., 1999), improved cognitive performance (Bruce-Keller *et al*., 1999), and lessened age-related cognitive deficits (Halagappa et al., 2007). The mechanism of neuroprotection by IF may be related to anti-inflammatory and antioxidant effects, improvement of hippocampal synaptic plasticity, and increased activity of neurotrophic factors but remain exclusive (Liu *et al*., 2019).

The gut microbiome modulates host brain function, including cognitive behavior, via the microbiota-gut-brain axis (Hu et al., 2016). Human and animal studies have revealed the vital activity of microbial metabolites generated by gut microbes in regulating central nervous system (CNS) disorders (Jiang et al., 2017) and in the pathology of AD and other neurodegenerative diseases (Giau et al., 2018). We showed that IF prevented synaptic structure damage and cognitive decline in mice with type 2 diabetes; prevention resulted from enhanced mitochondrial biogenesis and energy metabolism gene expression in the hippocampus, reshaping the gut microbiome (e.g., *Lactobacillus*, *Facklamia*, *Corynebacterium*, *Allobaculum*), and elevating the amounts of beneficial microbial metabolites, such as short-chain fatty acids and 3-indolepropionic acid (IPA) (Liu *et al*., 2020). Similarly, IF promoted nerve regeneration and recovery after chronic constriction injury, and recovery was associated with an increase in serum IPA (Giovanni et al., 2020). However, study aiming for exploring whether IF-induced alterations in the gut microbiome and microbial metabolites could prevent the cognitive deficits in AD there is surprisingly lacking.

In this study, we used APPswePSEN1dE9 (APP/PS1) mice, which accumulate Aβ plaque at the age of 7 months, to investigate the effects of IF on alleviating cognitive deficits in AD. Integrative modeling on hippocampus transcriptome, gut microbiome, and metabolome was conducted to uncover the biological mechanisms underlying the observed benefits of IF on alleviating cognitive impairments. Our findings suggest that the gut microbiota and the microbial metabolites have important functions in mediating the neuroprotective effects of IF in an AD animal model.

## Results

### IF alleviates cognitive impairment and amyloid deposition in a mouse model of AD

Four-month-old APP/PS1 mice (also known as AD mice) and wild-type (WT) littermates were treated either *ad libitum* or with an IF regimen at 24 h intervals for 16 weeks (Figure 1A). We recorded the weight of mice in each group on the fasting day (Figure 1B). The body weight change of AD mice was greater than the change for WT mice (Figure 1C; p<0.01). IF reduced the body weight gain of both genotypes of mice (Figure 1C p<0.01). Consistently, the average food intake every two days was greater for the AD mice compared with WT mice (Figure 1D; p<0.01). IF reduced the food intake of the AD mice (Figure 1D; p<0.01). Water intake by the AD mice was not different from that of the WT mice (Figure 1E). However, IF increased water intake by both genotypes of mice (Figure 1E; p<0.01). There was no significant sex difference in body weight change, food intake, or water intake after IF of AD mice (Figures S1A-S1C).The effect of IF on cognitive function was determined by Morris water maze test. The escape latency on the navigation tests was higher for the AD mice compared with the WT mice, which indicated that the AD mice had cognitive deficits (Figure 1F; p<0.01). IF decreased the escape latency in the day 6 navigation test, which indicated that IF improved cognition of the AD mice (Figure 1F; p<0.05). Interestingly, we found that IF improved the cognitive function of the male AD mice but not the female AD mice (Figure S1D). As shown in Figure 1G and Figure S1E, the trajectory of mice on the probe day was also recorded.

**Figure 1.**
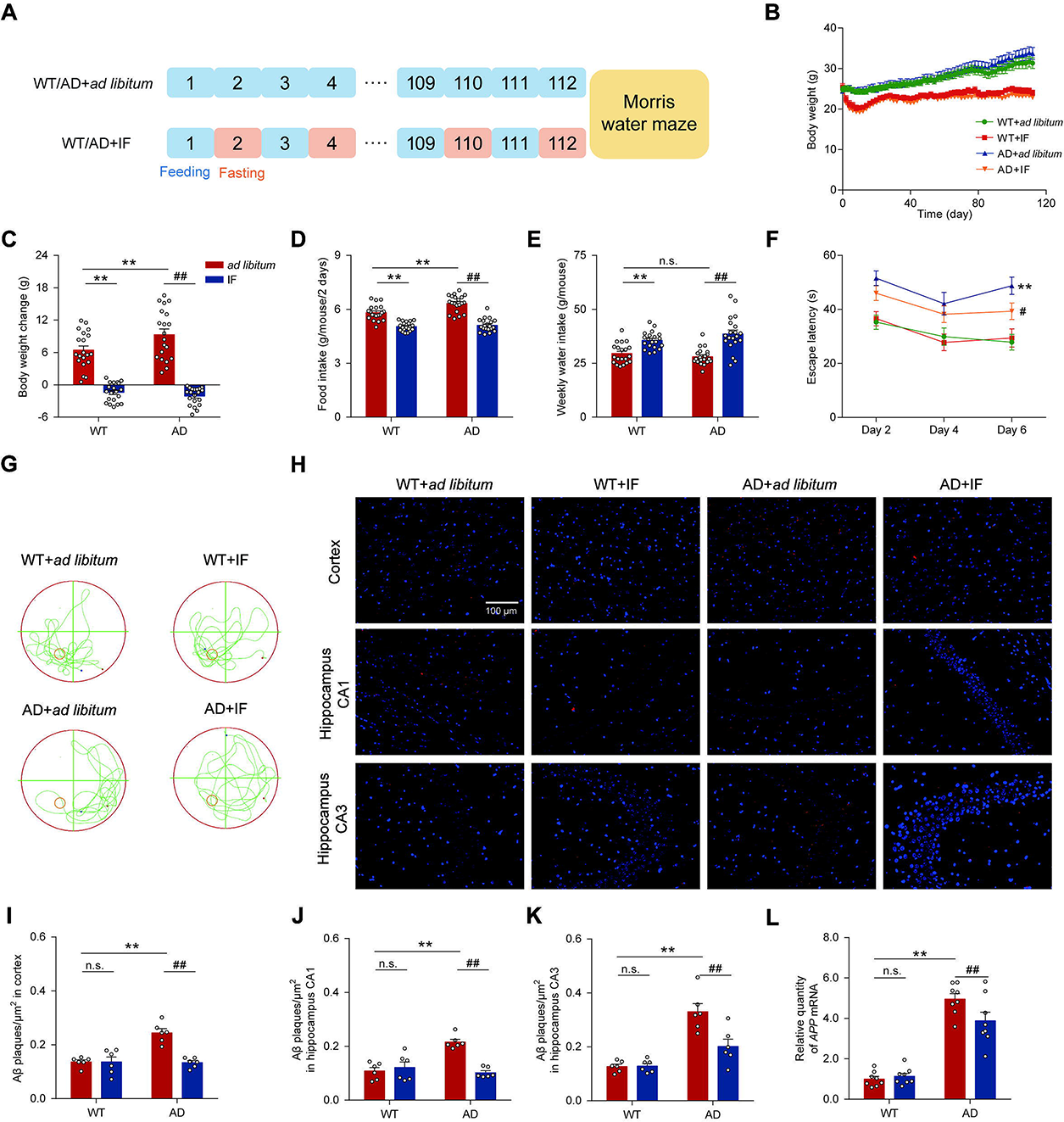
IF mitigated cognitive impairment of AD mice. (**A**) Schematic of the treatment with IF or *ad libitum* in each group. **(B)** Body weight trend. **(C)** Body weight change. **(D)** Food intake. **(E)** Weekly water intake. Cognitive function was measured by the Morris water maze test (see Methods); **(F)** Escape latency and **(G)** Mouse tracks during the navigation test (A-G, n=20, 10 male and 10 female mice per group) were recorded. **(H)** Representative immunofluorescence images of Aβ plaques in the cortex, CA1, and CA3 region of the hippocampus (n=6 slices, 3 male and 3 female mice per group). Scale bars: 100 µm. **(I-K)** Aβ plaque density (number of plaques/μm^2^) in the cortex, CA1, and CA3 region of the hippocampus (n=6, 3 male and 3 female mice per group). **(L)** mRNA levels of *APP* in the hippocampus (n=8, 4 male and 4 female mice per group). Data are mean ± SEM. *p < 0.05, **p < 0.01, compared with the WT+*ad libitum* group, ^#^p < 0.05, ^##^p < 0.01 compared with the AD+*ad libitum* group. Significant differences between means were determined by two-way ANOVA with Newman-Keuls multiple comparisons test. See also Figures S1.

The accumulation of Aβ plaques in the brains of AD mice was measured by immunofluorescence staining (Figure 1H and Figure S1F). IF reduced the Aβ plaque deposition in the cortex, CA1, and CA3 region of the AD mouse brain (Figures 1I-1K; p<0.01). There was no significant sex difference for Aβ plaque deposition in the cerebral cortex, hippocampal CA1, and hippocampal CA3 after IF of AD mice (Figures S1G-S1I). In the hippocampus of the AD mice, the mRNA of *APP*, the transgenic human Aβ precursor protein gene, was also downregulated by the IF regimen (Figure 1L; p<0.01). Note that IF downregulated *APP* expression in male AD mice but not in female AD mice. (Figure S1J).

### IF improves synapse ultrastructure and alters the expression of hippocampal genes in AD mice

The pathological features of AD are related to abnormal synaptic plasticity, which can be affected by the accumulation of Aβ plaques (Sun et al., 2019). Postsynaptic density (PSD) is a complex subcellular domain that forms an essential protein network and modulates synaptic plasticity (Moraes et al., 2021). We found that both the length and width of the PSD were shorter in the AD mice compared with the WT mice (Figures 2A-2C, Figure S2A; p<0.01). However, IF increased the length and width of the PSD in the AD mice hippocampus (Figures 2B and 2C, Figures S2B and S2C; p<0.01). Consistently, the mRNA of *PSD95*, an essential gene that regulates the structure of the PSD, was also upregulated in the AD mice hippocampus (Figure 2D, Figure S2D; p<0.05).

**Figure 2.**
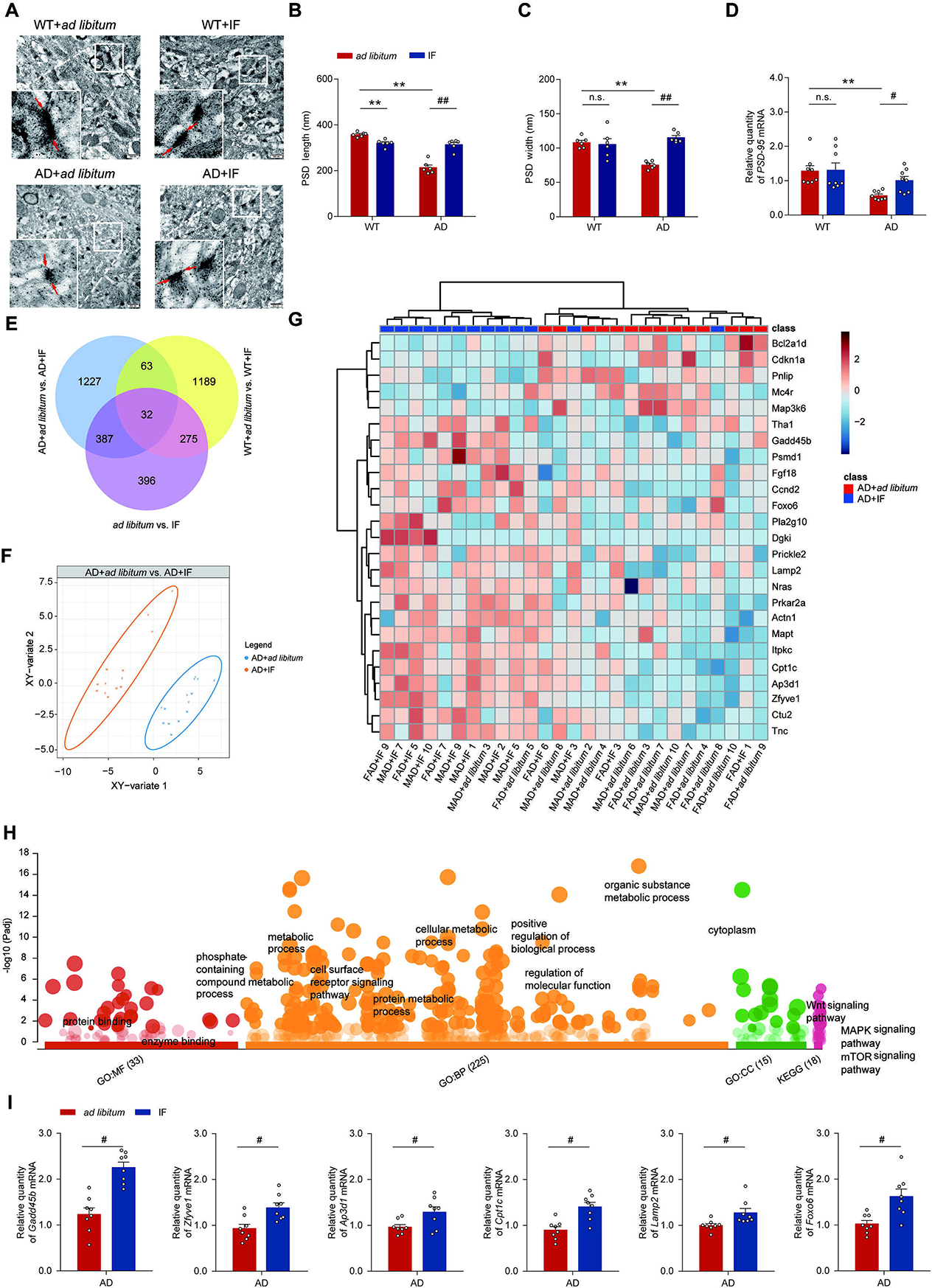
IF improved synapse ultrastructure and altered expression of hippocampal genes in AD mice. **(A)** Representative images of the ultrastructure of synapse in the hippocampus (n=6 slices, 3 male and 3 female mice per group). **(B-C)** The length and width of PSD (n=6, 3 male and 3 female mice per group). **(D)** *PSD-95* mRNA level in the hippocampus (n=8, 4 male and 4 female mice per group). **(E)** IF-induced alterations in genes expressed in the hippocampus of AD, WT, or in a combined group of both AD and WT mice without considering gene type (p<0.05). **(F)** GO and KEGG pathway annotations for 175 genes whose expression differed between AD+*ad libitum* and AD+IF. **(G)** A score plot revealing clear discrimination between AD+*ad libitum* vs. AD+IF. The plot was obtained by using the partial least squares regression-discrimination analysis on a selection of 175 genes expressed in the hippocampus. **(H)** A Z-score scaled heatmap of 175 differentially expressed genes between AD+*ad libitum* and AD+IF with p < 0.05. The top 25 genes are presented. **(I)** mRNA levels of *Gadd45b*, *Zfyve1*, *Ap3d1*, *Cpt1c*, *Lamp2*, and *Foxo6* in the hippocampus (n=8, 4 male and 4 female mice per group). Data are mean ± SEM. *p < 0.05, **p < 0.01, compared with the WT+ad libitum group, ^#^p < 0.05, ^##^p < 0.01 compared with the AD+*ad libitum*. Significant differences between mean values were determined by two-way ANOVA with Newman-Keuls multiple comparisons test. See also Figures S2.

We performed RNA sequencing with mice hippocampi to identify key biological processes and pathways that might be altered by IF. A total of 56,090 genes was detected, including 14,100 newly predicted genes lacking annotation. We found that 1709, 1559, and 1090 genes were differentially expressed in the hippocampus in AD mice, WT mice, and mice in a combined group that contained both AD and WT mice without considering genotype after IF (ANOVA, p<0.05). A large discrepancy existed in the panels of genes that differed significantly between AD+*ad libitum* and AD+IF, WT+*ad libitum* and WT+IF, and *ad libitum* and IF without considering genotype (Figure 2E). This discrepancy suggested a gene type effect on hippocampal response to IF regimen.

We focused on genes that were up- (n=1,103) or down- (n=606) regulated by IF in AD mice (p<0.05). By comparing IF to *ad libitum*, we found that differentially expressed genes in AD mice were enriched for 280 KEGG pathways (Table S1). Genes that were highly expressed after IF were enriched in Wnt signaling, mTOR signaling, autophagy, hippo signaling, Notch signaling, MAPK signaling, and sphingolipid signaling (p<0.05). Genes downregulated by IF were enriched in pathways related to primary bile acid and unsaturated fatty acids biosynthesis (p<0.05). Particular focus was given to 175 differentially expressed genes in AD+*ad libitum vs.* AD+IF mice proposed to be involved in pathways related to AD physiology, oxidative phosphorylation, neurotrophin signaling pathway, and synapse (Figure 2F, Table S2). We observed a clear discrimination between AD+*ad libitum* vs. AD+IF (Figure 2G). Expression of *Gadd45b*, *Zfyve1*, *Ap3d1*, *Cpt1c*, *Lamp2*, and *Foxo6* in AD mice hippocampi was greatly enhanced by IF (Figure 2H; Student’s t-test, p < 0.05); expression levels were confirmed by qPCR analysis (Figure 2I).

Like the results from the RNA sequencing, IF reduced the overactivation of microglia in the cerebral cortex and CA1 of hippocampal area of AD mice (Figures S2E-S2G; p<0.05). Moreover, IF downregulated the expression of TNF-α and upregulated the expression of IL-10 (Figures S2H and S2I; p<0.05).

### IF improves the gut barrier and restructured the gut microbiota of AD mice

The gut microbiome has vital functions in AD physiology (MahmoudianDehkordi et al., 2019). Disorders associated with gut microbiota can exacerbate inflammation and gut leakage, increase neuronal dysfunction and apoptosis, and consequently promote learning and memory impairment. As shown in Figure 3A and Figure S3A, compared with the WT group, there was severe mucosal myometrial injury, large erosion areas, and more inflammatory cell infiltration in the colon of the AD mice. IF increased crypt length (Figure 3B) and muscle layer thickness (Figure 3C) of the AD mice (Two-way ANOVA with Newman-Keuls test, p<0.01). To further assess the integrity of the gut barrier, the expression of claudin-1, an essential tight junction protein of the gut barrier, was measured by immunohistochemical staining. IF increased expression of colon claudin-1 in the AD mice (Figure 3D; Two-way ANOVA with Newman-Keuls test, p<0.01). There was no significant sex difference in the effect of IF intervention on either crypt length, the thickness of the muscle layer, or claudin-1 protein expression in the colon of AD mice (Figures S3B-S3D).

**Figure 3.**
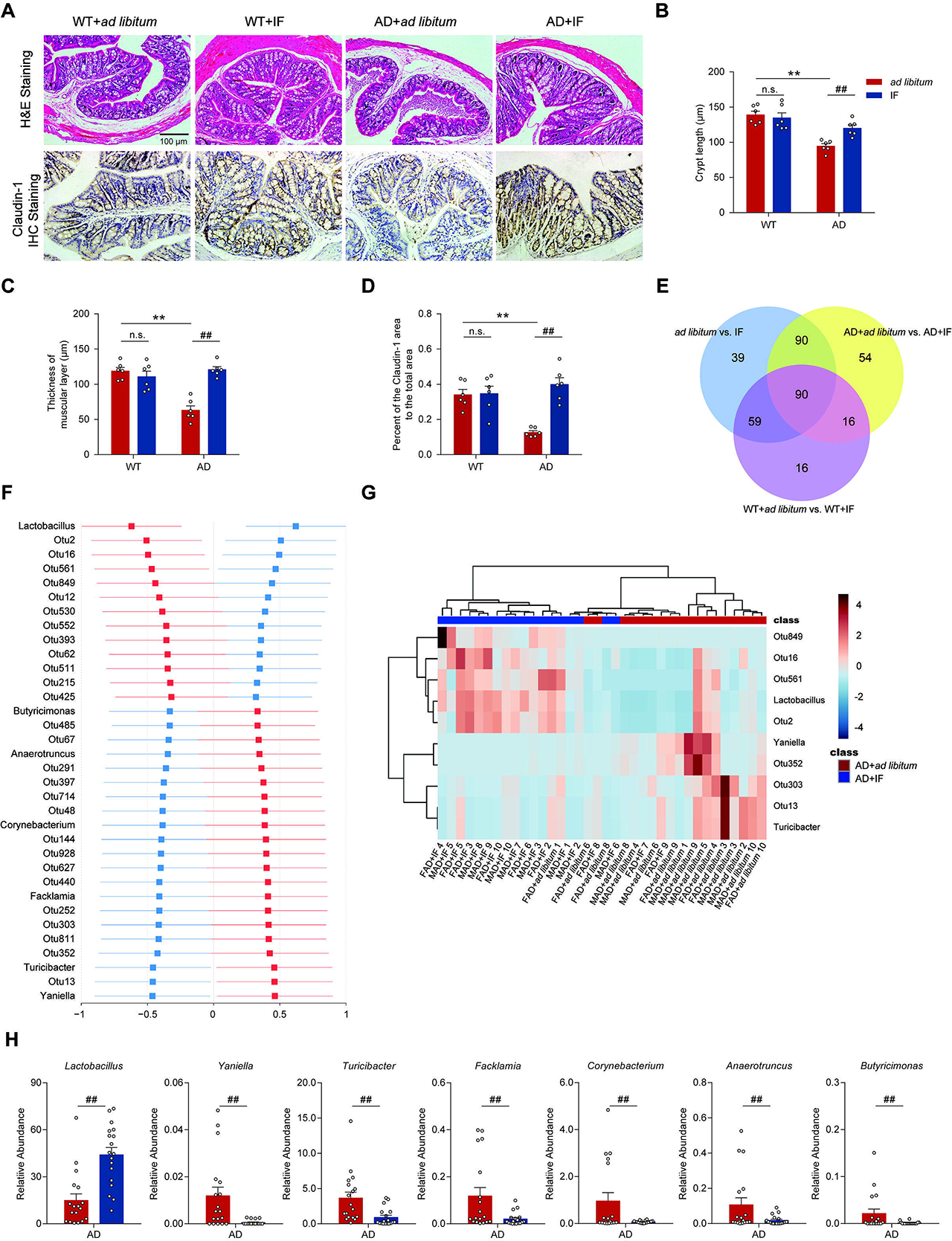
IF improved the gut barrier and restructured the gut microbiota of AD mice. **(A)** Representative images of hematoxylin and eosin (H&E) staining and claudin-1 protein immunohistochemistry (IHC) of the colon (n=6 slices, 3 male and 3 female mice per group). **(B)** Crypt length (n=6, 3 male and 3 female mice per group). **(C)** The thickness of the muscular layer (n=6, 3 male and 3 female mice per group). **(D)** Quantitative immunohistochemical analysis of claudin-1 protein in colon (n=6, 3 male and 3 female mice per group). **(E)** IF-induced alterations in gut microbiome in AD, WT, and in a combined group containing both AD and WT mice without considering gene type. Optimal panels of bacteria that discriminated between *ad libitum* and IF with or without considering gene type were determined from 52 genera and 430 OTUs by using partial least square discriminant analysis incorporated into a repeated double cross-validation framework with an all-relevant variable selection procedure. **(F)** Differences in a selection of six genera and 28 OTUs differed between AD+*ad libitum* and AD+IF group. The least-square means ± 95% confident intervals obtained from ANOVA (p<0.05) are presented. **(G)** A Z-score scaled heatmap of top 10 bacteria that differed between AD+*ad libitum* and AD+IF with p < 0.05. **(H)** The relative abundance of differential AD+*ad libitum* and AD+IF genera. Data are mean ± SEM. *p < 0.05, **p < 0.01, compared with the WT+*ad libitum* group, ^#^p < 0.05, ^##^p < 0.01 compared with the AD+*ad libitum* group. Significant differences between mean values were determined by two-way ANOVA with Newman-Keuls multiple comparisons test. See also Figures S3, S4.

Gut microbiota composition was determined from mouse fecal samples using 16S rRNA gene v3-v4 amplicon sequencing. IF did not have any significant effect on alpha and beta diversity of gut microbiota in the *ad libitum* group (Figures S3E and S3F), but alpha and beta diversity differed between *ad libitum* and IF groups in the relative abundance of microbial genera and OTUs (Figure 3E). Specifically, we constituted optimal panels of bacteria at genus and OTU levels to distinguish *ad libitum* and IF group, with or without considering genotype. The panels were derived by using partial least square-discriminant analysis incorporated into a repeated double cross-validation framework with an all-relevant variable selection procedure. In comparison with *ad libitum*, IF affected 250 bacterial species in the AD group (Figure 3E). Most of the differential bacteria were well-represented in the multivariate modeling-derived optimal panels of bacteria that discriminated WT+*ad libitum* and WT+IF, or *ad libitum* and IF including both AD and WT mice. This result indicated the robustness of IF-induced changes in the abundance of bacteria. When assessed individually, IF caused significant changes in relative abundance of six genera and 28 OTUs in AD mice (Figure 3F). As expected, these bacteria collectively contributed to a clear separation between *ad libitum* and AD+IF (Figure S4A). Compared with AD mice, IF greatly promoted the growth of *Lactobacillus* and several OTUs belonging to *Lactobacillus*, whereas led to a lower relative abundance of *Turicibacter* and *Yaniella* (Figures 3G and 3H). A PICRUSt analysis revealed that genes upregulated by IF were enriched in pathways of apoptosis, the metabolism of primary and secondary bile acids, and fatty acid biosynthesis, compared with the *ad libitum* group of AD mice (Table S3).

### Effects of IF on plasma metabolites in AD mice

IF greatly affected the plasma metabolome, and notably, a gene-type specific effect on IF-induced changes in plasma metabolites was observed Specifically, 348 and 182 metabolites were optimally selected as key metabolites relevant to IF in the AD and WT group, respectively, with 77 metabolites in common. Univariate statistics revealed 87 differential AD+*ad libitum vs.* AD+IF metabolites (Figure S5A; ANOVA, p<0.05) showing a varying degree correlations with each other (Figure S5B). These metabolites were enriched in the metabolism of histidine, beta-alanine, arginine, and proline (Figure 4C; p<0.05). Of note, 12 metabolites have been associated with AD development and/or cognitive impairment (Figure 4D; p<0.05). Metabolites involved in the tryptophan metabolic pathway have been considered gut microbiota-related metabolites, including IPA and indolelactic acid.

**Figure 4.**
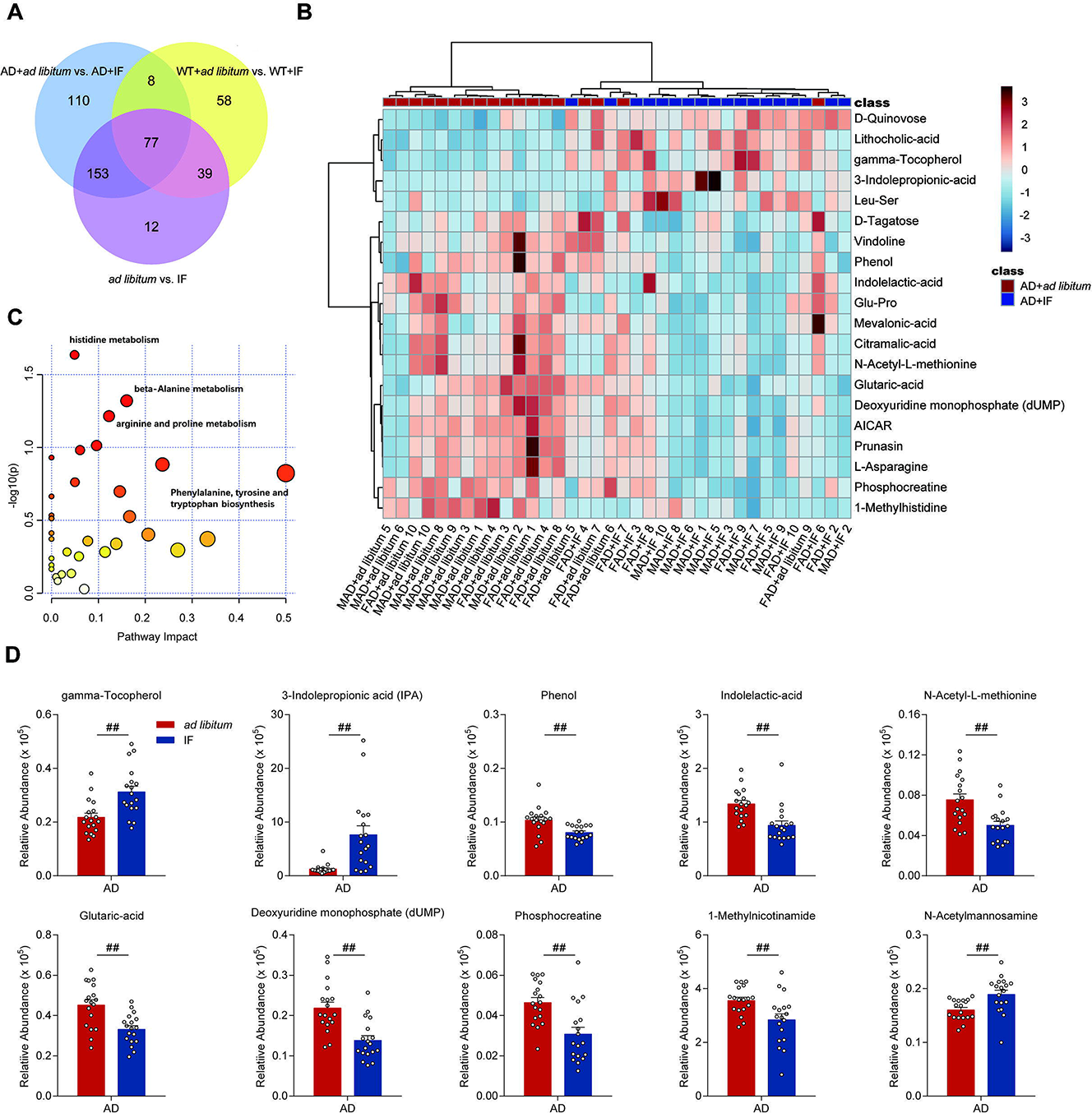
IF affected plasma metabolome of AD mice. (**A**) IF induced alterations in AD, WT, and in a combined group containing both AD and WT mice without considering gene type. Optimal panels of metabolites that discriminated between *ad libitum* and IF with or without considering gene type were determined by using partial least square-discriminant analysis incorporated into a repeated double cross-validation framework with an all-relevant variable selection procedure. (**B**) A Z-score scaled heatmap of differential metabolites between AD+*ad libitum* and AD+IF with p < 0.05. The top 20 differential metabolites are presented. (**C**) Metabolic pathways influenced by IF treatment in AD mice. (**D**) Plasma levels of gamma-tocopherol, X3-Indolepropionic acid, phenol, indolelactic-acid, N-acetyl-L-methionine, glutaric-acid, deoxyuridine monophosphate, phosphocreatine, 1-methylnicotinamide, and N-acetylmannosamine (n=18, 9 male and 9 female mice per group). See also Figures S5.

### Multi-OMICs integration for IF

Having identified hippocampus genes (n=175), gut bacteria (n=34), and plasma metabolites (n=87) that were key signatures relevant to IF treatment of AD mice, we then measured the interplay of these features to assess potential mechanistic connections (Figure 4). We first confirmed the validity of genes, gut bacteria, and metabolites in discriminating AD+IF from AD+*ad libitum*, with an accuracy of 92%, 100%, and 92%, respectively (Figure 5A). This prediction was performed by a partial least square-discriminant analysis incorporated into a repeated double cross-validation framework, a widely used strategy that minimizes the risk of statistical overfitting. The predictive ability of the models outperformed 1000 permuted models, which demonstrated the robustness and validity of predictive models with great generalizability (Student’s t-test, p < 0.05). We then performed the DIABLO integration analysis using a weighted design (Figure S6A) to identify biologically relevant and highly correlated OMICs signatures (Figure S6B). DIABLO analysis identified one latent component composed of 28 genes, 32 bacteria, and 40 metabolites as key OMICs signatures that contributed to separation between AD+*ad libitum* and AD+IF (Figures 5B and 5C). The selected signatures were highly correlated (Figure 5D). Furthermore, despite the inter-relationship of differential AD+*ad libitum vs.* AD+IF genes, bacteria, and plasma metabolites, we also found strong correlations between integrated key predictors and IF-induced improvements in escape latency on the navigation tests (r=0.51, 0.50, and 0.45 for genes, gut bacteria, and plasma metabolites, respectively).

**Figure 5.**
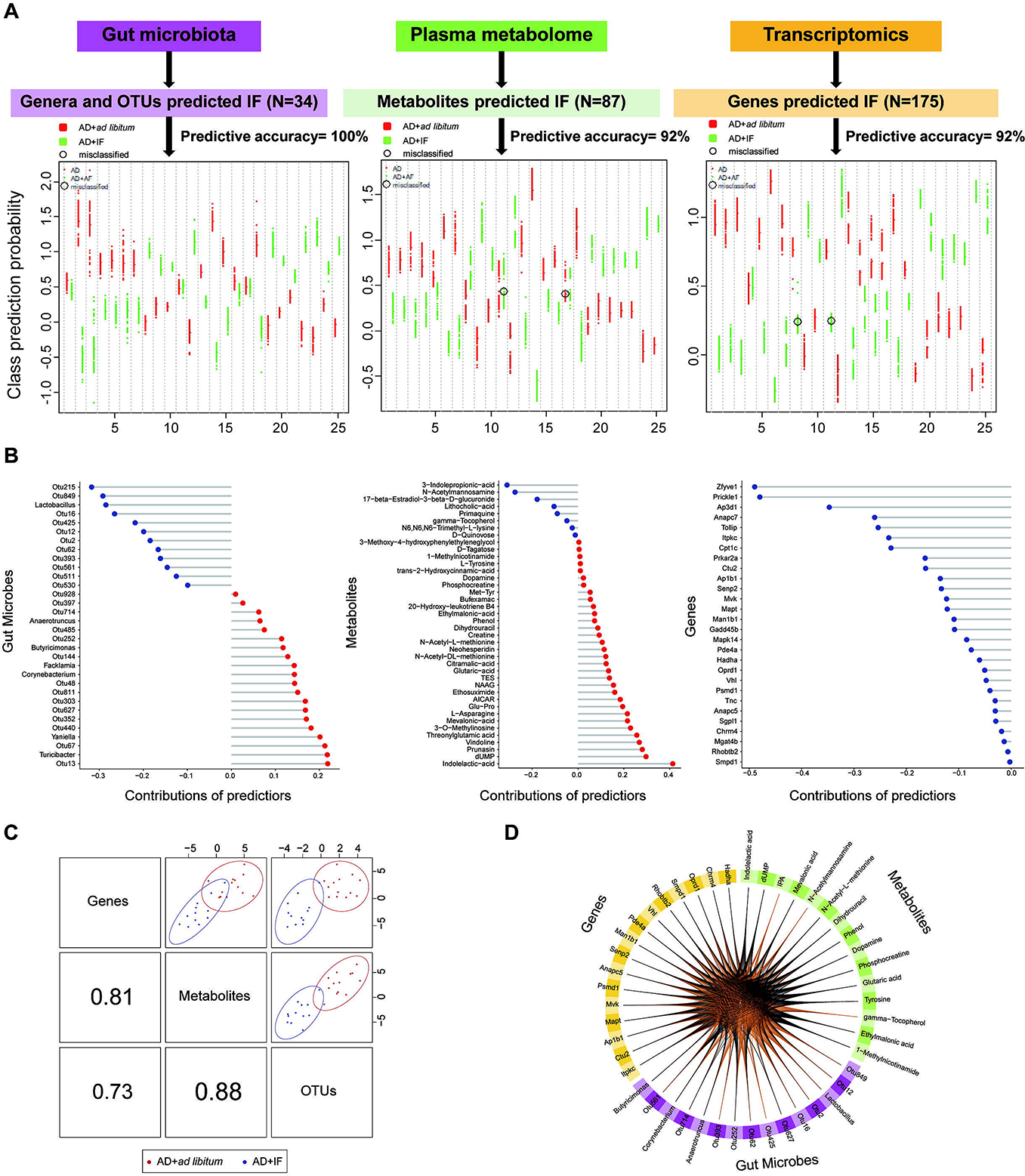
Multi-OMICS integration for IF treatment. **(A)** The performance of predictive models for signatures of IF status. OMICs signatures included 175 differential AD+*ad libitum* vs. AD+IF genes expressed in the hippocampus, 34 gut bacteria, and 87 plasma metabolites. For each data set, multivariate predictive modeling was conducted using partial least square-discriminant analysis incorporated into a repeated double cross-validation framework. Prediction performance is shown in downstream figures: each swim lane represents one mouse. For each sample, class probabilities were computed from 200 double cross-validations. Class probabilities are color-coded by class and presented per repetition (smaller dots), and averaged over all repetitions (larger dots). Misclassified samples are circled. Predictive accuracy was calculated as several correctly predicted samples/total number of measured samples. **(B)** Lollipop plot of contributions of key OMICs signatures identified by DIABLO integrative modeling for discriminating AD+IF from AD+*ad libitum*. Loading of DIABLO integrative modeling for each predictor is presented. Blue bars indicate IF-induced improvements in predictors. Predictors that were lower in AD+IF compared with AD+*ad libitum* are in red. **(C)** Model performance of DIABLO integrative analysis of OMICs signatures concerning IF. The use of DIABLO maximized the correlated information between genes, bacteria, and metabolites. Scatter plots depicting the clustering of groups, i.e., AD+*ad libitum* and AD+IF, based on the first component of each data set from the model, showed significant separation between groups. A scatterplot displays the first component in each data set (upper diagonal plot) and Pearson correlation between components (lower diagonal plot). **(D)** The Circos plot shows the positive (negative) correlation, denoted as brown (gray) lines, between selected multi-omics features. See also Figures S6.

### The neuroprotective effects of IF are mediated by gut microbiota

Ten-month-old APP/PS1 and WT littermates were administered with an antibiotic mixture (ABx; see Methods) for two weeks and then treated with IF for six weeks (Figure 6A). It showed that ABx treatment significantly reduced the 16S rRNA copies in feces(Figure S7A). During the experiment, the body weight, food intake, and weekly water intake of the mice were recorded (Figures S7B-S7E). IF reduced the body weight and food intake of AD mice but did not affect water intake (Figures S7B-S7E; p<0.05). Compared with the AD+IF group, the AD+IF+ABx group did not have significant changes in body weight, food intake, and weekly water intake (Figures S7B-S7E).

**Figure 6.**
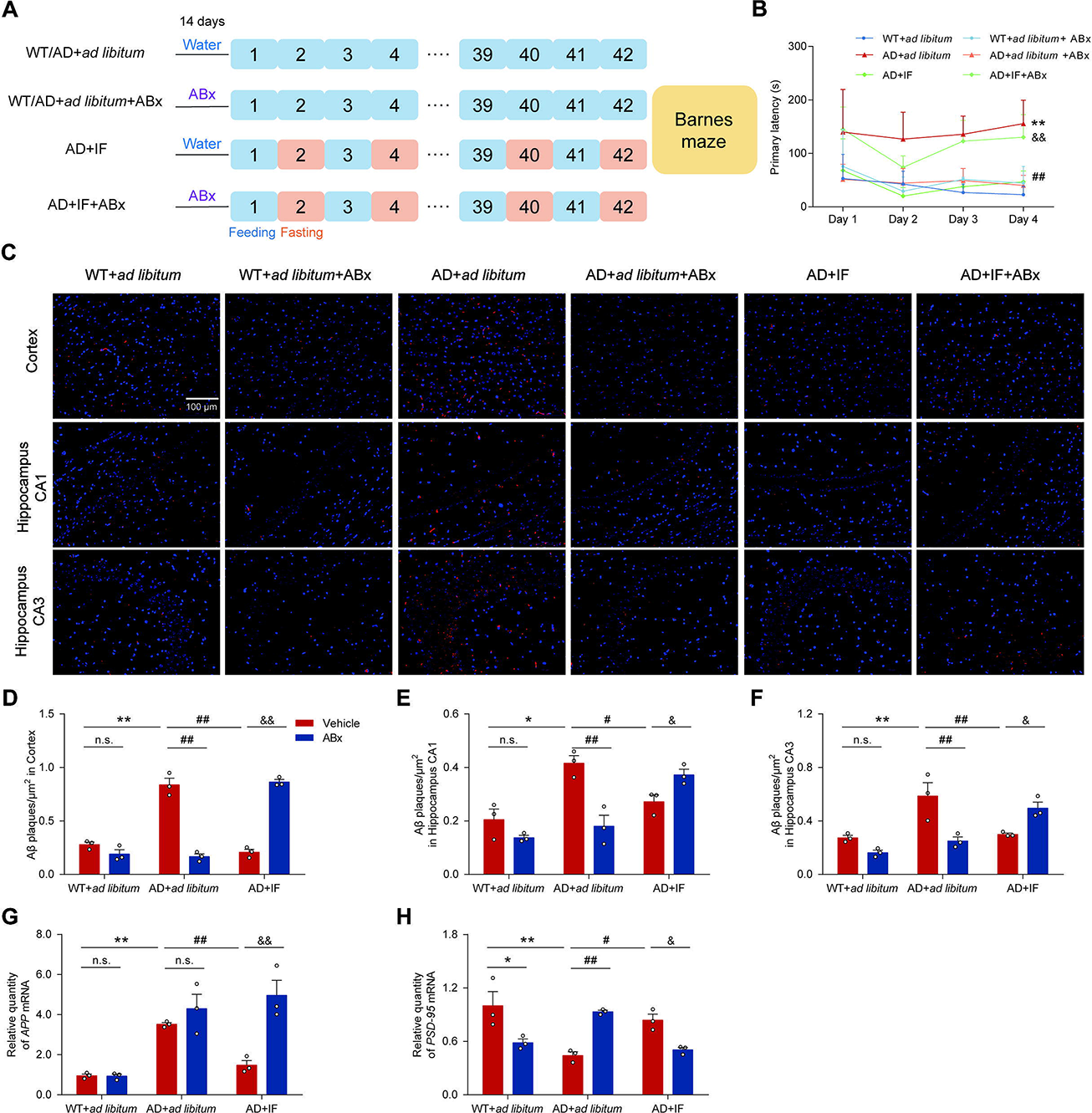
Improvement of cognitive function in the AD mice by IF was mediated by gut microbiota. **(A)** Schematic of the treatment with ABx and IF in each group. **(B)** Primary latency in acquisition trials of Barnes maze (WT+*ad libitum*, WT+*ad libitum*+ABx, AD+*ad libitum*+ABx, AD+IF: n=5; AD+*ad libitum*, AD+IF+ABx: n=4). **(C)** Representative images of immunofluorescence staining of Aβ plaques in the cortex, CA1, and CA3 region of the hippocampus (n=3 male mice per group). Scale bars: 100 µm. **(D-F)** Aβ plaque density (number of plaques/μm^2^) in the cortex, CA1, and CA3 region of the hippocampus (n=3 male mice per group). **(G)** The mRNA levels of *APP* in the hippocampus (n=3 male mice per group). **(H)** The mRNA levels of *PSD-95* in the hippocampus (n=3 male mice per group). Data are mean ± SEM. *p < 0.05, **p < 0.01, compared with the WT+*ad libitum* group, ^#^p < 0.05, ^##^p < 0.01 compared with the AD+*ad libitum* group, and ^&^p < 0.05, ^&&^p < 0.01 compared with the AD+IF group. Significant differences between mean values were determined by two-way ANOVA with Newman-Keuls multiple comparisons test. See also Figures S7.

To test spatial memory ability, the Barnes maze test was conducted. In acquisition trials and the probe trial, with increasing training days, compared with the WT+*ad libitum* group, the primary latency of the AD+*ad libitum* group increased (Figure 6B and Figure S7F; p<0.01). IF improved the cognitive function of AD mice. However, compared with the AD+IF group, the primary latency of AD+IF+ABx was significantly increased, which indicated that the improvement in cognitive function of AD mice by IF was partly offset by the removal of gut microbiota (Figure 6B and Figure S7F; p<0.01). In addition, we recorded the number of times each mouse poked its head into each hole (Figure S7G).

The improvement effect of IF on Aβ deposition was eliminated by ABx treatment (Figures 6C-6F). IF significantly downregulated the expression of *APP* mRNA in the hippocampus of AD mice (Figure 6G; p<0.01). However, IF had no significant effect on *APP* mRNA expression in the hippocampus of AD mice treated with ABx (Figure 6G). In addition, IF significantly upregulated the expression of *PSD-95 and IL-10* mRNA and upregulated the expression of *TNF-α* mRNA in the hippocampus of AD mice (Figure 6H and Figures S7H and S7I). However, IF-induced upregulation of *PSD-95 and IL-10* expression in AD mice was offset by the removal of gut microbiota.

## Discussion

Many studies have proved the potential of an IF regimen as an effective nutritional intervention for improving cognitive function (Hu *et al*., 2017; Liu *et al*., 2019; Liu *et al*., 2020; Ooi *et al*., 2020; Vasconcelos *et al*., 2014). We found that IF improved cognitive function and spatial memory and attenuated the Aβ accumulation in the cortex and hippocampus of APP/PS1 mice. Notably, IF beneficially altered autophagy-, synaptic plasticity-, and cognitive function-related gene expression in the hippocampus of AD mice. Similarly, IF improved the gut barrier structure, reshaped the gut microbiome, and improved the production of neuroprotective microbial metabolites. Pseudo germ-free mice were employed to further demonstrate a pivotal function of gut microbiota in regulating the neuroprotective effects of IF in AD mice. Benefiting from an integrated analysis of gene expression, gut microbiome, and metabolites, our findings indicated that 16 weeks of IF alleviated cognitive deficits in APP/PS1 mice, predominately via a gut-brain axis.

In addition to genetic factors, AD development is influenced by environmental factors such as diet (Silva *et al*., 2019). Previous studies showed that IF improved spatial learning and memory in various transgenic AD mouse models, *e.g.*, APP mutant mice, 3xTgAD mice, and APP/PS1 mice (Halagappa *et al*., 2007; Lazic et al., 2020; Liu *et al*., 2019; Zhang *et al*., 2017). As measured by behavioral tests, we observed a significant improvement in the cognitive function of APP/PS1 mice after IF treatment (Figure 1). Although the “amyloid cascade hypothesis” has encountered some resistance and controversy, Aβ has been consistently related to the occurrence and development of AD (Li et al., 2008). Moreover, elevated β-amyloid and tau are considered important biomarkers/predictors for early dementia (Hammond et al., 2020). Herein, we found that IF lessened the excessive accumulation of Aβ in the cerebral cortex and hippocampal CA1 and CA3 regions of AD mice and greatly downregulated the expression of *APP* in the hippocampal region (Figure 1). IF reduced Aβ deposition in the brains of AD mice, which indicated the benefits of IF in AD prevention.

Accumulation of Aβ in the brain is caused by an imbalance in the production and clearance of Aβ (Wani et al., 2019). Autophagy is an important mechanism of Aβ clearing (Wani *et al*., 2019). Our study showed that IF increased autophagy-related gene expression, including expression of *ZFYVE1*, *Lamp2*, and *Foxo6,* in AD mice hippocampus (Figure 2). This finding may partly explain the beneficial effects of IF on Aβ deposition. *ZFYVE1*/*DFCP1* (zinc finger, FYVE domain containing 1) is a phosphatidylinositol-3-phosphate-binding protein that promotes the nucleation step of autophagosome formation (Godar et al., 2015). A recent study indicated that *Lamp2* (lysosome-associated membrane protein 2)-deficient rats had cognitive and locomotion dysfunction (Ma et al., 2018). FoxO6, highly enriched in the adult hippocampus, promotes memory consolidation by regulating a program coordinating neuronal connectivity in the hippocampus (Salih et al., 2012). In addition, IF upregulated the expression of *Gadd45b*, a regulator of neurogenesis, and *Ap3d1* and *Cpt1c* (Figure 2). Knockout of *Cpt1c* in CNS impaired spatial memory in an animal model, which could be explained by the function of *Cpt1c* in regulating the level of ceramide in the endoplasmic reticulum of hippocampal neurons (Carrasco et al., 2012; Luigi Pinto, 2014). Moreover, we found that IF significantly protected the ultrastructure of the synapse in hippocampal neurons and elevated the expression of synaptic plasticity-related genes including *PSD-95* and *Ap3d1*. *Ap3d1* is related to synaptic plasticity (Lachén-Montes et al., 2017) and AD phenotype (Zhang et al., 2018). Synaptic activity interruption and synaptic loss are early events in AD pathogenesis that precede the formation of the cerebral cortex, protein deposition in the brain, or clinical manifestations of the disease (Sheng et al., 2012).

In AD, neuroinflammation, instead of being a mere bystander activated by emerging senile plaques and neurofibrillary tangles, contributes to the pathogenesis, as do the plaques and tangles themselves (Zhang et al., 2013a). In this study, IF reduced the activated microglia, downregulated expression of the inflammatory mediator TNF-α, and upregulated anti-inflammatory cytokine IL-10 (Figure S2). In agreement with our results, a recent human study revealed that a 6-month alternate-day fasting regimen reduced the serum level of age-associated inflammatory markers (Stekovic et al., 2019). Also, IF prevented neuroinflammation and cognitive deficits by stimulating the adaptive responses in the CNS and peripheral system in a rat model of sepsis (Vasconcelos *et al*., 2014). However, in a 5xFAD transgenic mice model, alternate-day fasting exacerbated neuroinflammation and enhanced the protein level of TNF-α (Lazic *et al*., 2020). IF also exacerbated the acute immune and behavioral sickness response by increasing inflammatory mediators TNF-α and IL-6 in a viral poly(I:C)-mimic mouse model (Zenz et al., 2019). Our recent work proved that, compared with time-restricted feeding and intermittent energy restriction, alternate-day fasting exacerbated the DSS-induced colitis and inflammatory responses in a chronic colitis mouse model (Zhang et al., 2020). These conflicting findings indicate that more research is needed to explain the differences in IF regimen-regulating effects in different animal models and human clinical trials.

Gut dysbiosis and gut barrier leak are associated with the neuroinflammatory responses in CNS (Köhler et al., 2016). Thirty percent caloric restriction (CR) prevented Aβ accumulation in Tg2576 AD mice, partly by modulating gut microbiota composition (Cox et al., 2019). We found that IF greatly influenced the relative abundance of bacteria related to cognitive function, such as *Lactobacillus*, *Corynebacterium*, and *Allobaculum* (Figure 3 and Figure S3). *Lactobacillus* was shown to mediate the effects of long-term or short-term CR on alleviating the systemic inflammatory responses and improving energy metabolism (Pan et al., 2018; Zhang et al., 2013b). Supplementation with *Lactobacillus* attenuated the cognitive dysfunction and anxiety-like behaviors in inflammatory bowel disease via the gut-brain-microbiota axis (Emge et al., 2016; Smith et al., 2014). Moreover, oral administration of probiotics containing *Lactobacillus reuteri* improved spatial memory, reduced Aβ plaques, and reduced IL-1β and TNF-α as inflammation markers in AD rats with Aβ_1-40_ intra-hippocampal injection (Mehrabadi and Sadr, 2020). Supplementation with *L. reuteri* also improved cognitive function and lipid peroxidation and lessened neuronal damage in a PD rat model (Alipour Nosrani et al., 2021). Conversely, IF downregulated the level of *Butyricimonas* and *Allobaculum* (Figure 3). Recent research indicated that the relative abundance of *Butyricimonas* was negatively correlated with human cognition ability (Ren et al., 2020). In our previous research, *Allobaculum* was reduced in IF-treated diabetic mice (Liu *et al*., 2020).

The IF regimen had a great effect on the plasma metabolome, particularly for metabolites associated with AD development and/or cognitive impairment. Notably, a multi-omics integrative analysis revealed strong connections between IF-induced alterations in the expression of hippocampus genes related to autophagy, synaptic plasticity, and inflammatory responses and relative abundances of gut microbiota and serum metabolites related to AD phenotype. These findings suggest a mediating activity of the gut-brain axis in the benefits of IF for AD (Lachén-Montes *et al*., 2017; Tuomainen et al., 2018). In agreement with our results, *Lactobacillus reuteri* prevented the intestinal epithelial toxicity in a lipopolysaccharide-induced intestinal epithelial cell mouse model by enhancing antioxidant activities and tight junctions and attenuating apoptosis and autophagy via the mTOR signaling pathway (Han et al., 2016). IF led to an increase in serum IPA, a microbial tryptophan metabolite, in AD mice (Figure 4), in agreement with our previous finding that IF enhanced IPA generation in diabetic mice (Liu *et al*., 2020). Production of IPA is fully dependent on the presence of gut microbiota such as *Clostridium sporogenes* and *L. reuteri (Roager and Licht, 2018; Rothhammer et al., 2016)*. A recent study showed that IPA alleviated Aβ fibril formation (Chyan et al., 1999), and IPA has also been suggested as a promising candidate for the treatment of metabolic disorders by lowering fasting blood glucose and reducing insulin resistance (Abildgaard et al., 2018). IPA was found to reduce the levels of proinflammatory cytokines produced by lipopolysaccharide-activated astrocytes (Garcez et al., 2020). Moreover, IPA bound the pregnane X receptor (PXR) and suppressed inflammatory responses (Tuomainen *et al*., 2018), and, notably, knockout of PXR led to cognitive dysfunction and anxiety-like behavior (Boussadia et al., 2018). Thus, IPA might mediate the beneficial effects of IF of AD mice by activation of PXR. In addition, IF led to increases in gamma-tocopherol and N-acetylmannosamine in the serum of AD mice. These metabolites negatively correlated with Aβ accumulation (Morris et al., 2015; Yonekawa et al., 2014). Conversely, IF reduced serum levels of phosphocreatine and glutaric acid, two metabolites that were higher in AD patients than in healthy individuals (Rijpma et al., 2018). Further investigation is needed to explain the relationship between IF-induced changes in hippocampus gene expression and gut microbes and microbial metabolites.

The beneficial effects of IF on cognitive function, Aβ accumulation, and neuroinflammatory responses in AD mice were partly abolished by removing the gut microbiota (Figure 6 and Figure S7). Long-term broad-spectrum combinatorial antibiotic treatment was found to decrease Aβ plaque deposition (Minter et al., 2016). Similarly, our previous research also found that antibiotic treatment improved cognitive function in diabetic mice (Liu *et al*., 2020). The removal of gut microbiota by antibiotics abolished the positive effects of IF on improving cognition. These findings indicated that gut microbiota alteration mediates the beneficial effects of IF on the CNS.

In summary, the present study indicated that IF significantly improved the cognitive function of AD mice, which was accompanied by a decrease in Aβ accumulation and suppression of neuroinflammation in the CNS. The benefits of IF on cognitive deficits of AD might be attributed to an IF-improved relative abundance of *Lactobacillus*, enhanced generation of metabolites such as IPA, protection of the gut barrier, and prevention of neuroinflammation and Aβ accumulation in the CNS. These findings may suggest a therapeutic application and nutritional intervention for AD and other neurodegenerative diseases.

## Author contributions

M.J., Y.Z., L.S., X.H., J.Z., C.D., Y.S., X.L., X.J., X.D., Z.L., and L.S. performed the experiments and analyzed the data, Z.L., X.L., Y.Z., and M.J. designed the study, Z.L., L.S. Y.Z., and M.J. wrote the paper. M.J. and L.S. prepared the figures. All authors discussed the results and commented on the paper.

## Acknowledgments

**Funding:** This work was financially supported by the National Natural Science Foundation of China (Nos. 81871118 and 81803231), a General Financial Grant from China Postdoctoral Science Foundation (No. 2016M602867), a Special Financial Grant from China Postdoctoral Science Foundation (No. 2018T111104), and the Innovative Talent Promotion Program-Technology Innovation Team (2019TD-006). Dr. Zhigang Liu is also funded by the Tang Cornell-China Scholars Program from Cornell University in the U.S. and the Alexander von Humboldt-Stiftung in Germany.

The authors also thank AiMi Academic Services (www.aimieditor.com) for English language editing and review services.

## Data and code availability

All data needed to evaluate the conclusions in the paper are present in the paper and/or the Supplementary Materials. Additional data related to this paper may be requested from the authors.

The raw and processed data of RNA-sequencing in the current study were deposited on GEO (access is only available for reviewer and editor at present) (https://www.ncbi.nlm.nih.gov/geo/query/acc.cgi?acc=GSE164461), the accession number is **chityiimnngzxyr**. The accession number for the entire 16S rRNA sequencing dataset is available for reviewer by request. The other raw data were deposited in *figshare* dataset as follows: https://figshare.com/s/e73cf65b4258206f7b31

## STAR Methods

### Animals and treatment

Four-month-old APPswePSEN1dE9 (APP/PS1) double-transgenic mice (twenty male and twenty female) and forty age-matched wild-type littermates, on a B6C3-Tg background, were purchased from Model Animal Research Center of Nanjing University (Nanjing, China) and bred in the Northwest A&F University animal facility. Amyloid Precursor Protein (*APP*) and Presenilin1 (*PSEN1*) are regarded as causative genes for AD. Mice were housed singly with a 12-h light/dark cycle, at 22 ± 2°C and 50 ± 10% humidity in the Northwest A&F University animal facility. All mice were adaptively fed to 4 months of age. To identify mouse genotype, the DNA of each mouse was isolated from the ear and amplified by PCR with the following primers: APP: 5’-AGGACTGACCACTCGACCAG-3’ (forward) and 5’-CGGGGGTCTAGTTCTGCAT-3’ (reverse); PS1: 5’-AATAGAGAACGGCAGGAGCA-3’ (forward) and 5’-GCCATGAGGGCACTAATCAT-3’ (reverse); reference gene: 5’-CTAGGCCACAGAATTGAAAGATCT-3’ (forward) and 5’-GTAGGTGGAAATTCTAGCATCATCC-3’ (reverse). All animal experimental procedures were conducted following the guidelines for the care and use of animals, Eighth Edition (ISBN-10: 0-309-15396-4). The Northwest Agricultural and Forestry University of Science and Technology review committee approved the study (BGI-IRB).

All treatments were implemented on wild-type (WT) or APP/PS1 mice (male and female) randomly divided into *ad libitum* or IF diet groups, including WT+*ad libitum* (wild-type mice, n=20), WT+IF (wild-type mice with IF treatment, n=20), AD+*ad libitum* (APP/PS1 mice, n=20), AD+IF (APP/PS1 mice with IF treatment, n=20) for 16 weeks. The IF protocol comprised a day with enough food followed by a day of complete fasting. Food (AIN-93M, TROPHIC Animal Feed High-tech Co., Ltd. Nantong, China) was provided or removed every day at 9:00 am, with water supplied *ad libitum*. Mice were measured for body weight and food intake every two days and weekly for water intake. All animals performed behavioral tests and then were sacrificed to collect fecal samples, serum, and hippocampus tissue under liquid nitrogen and stored at −80°C for histological analysis and multi-omics study, 16S rRNA gene sequencing, plasma metabolome, and hippocampus transcriptome analyses, respectively. In addition, tissue samples of brain and colon from three mice in each group were collected and fixed, paraffin-embedded, and sectioned at 5 μm for pathology staining.

The set of animals in the validation test were divided into six groups: WT+*ad libitum* (n=5), WT+*ad libitum*+ABx (n=5), AD+*ad libitum* (n=4), AD+*ad libitum*+ABx (n=5), AD+IF (n=5), AD+IF+ABx (n=4). The IF regimen was the same as the aforementioned sets. Antibiotic cocktail (penicillin G sodium 1 g/L, metronidazole 1 g/L, neomycin sulfate 1 g/L, streptomycin sulfate 1 g/L, vancomycin hydrochloride 0.5 g/L) was given in the drinking water starting 14 days before the IF regimen and throughout the experiment (Figure 6A). The 16S rRNA transcripts in feces after antibiotic treatment were detected by qPCR (Figure S7A) as described (Wang et al., 2018). After six weeks of IF, the Barnes maze test was used to evaluate the spatial memory and learning ability of mice, and then biochemical indexes were assayed.

All of the experimental procedures were followed using the Guide for the Care and Use of Laboratory Animals: Eighth Edition (ISBN-10: 0-309-15396-4). We have complied with all relevant ethical regulations for animal testing, and research and protocols were approved by the Northwest A&F University, and BGI Institutional Review Board on Bioethics and Biosafety (BGI-IRB).

### Morris water maze test

The Morris water maze (MWM) test was performed to evaluate the capability of spatial learning and memory as described (Zhao et al., 2019). The diameter of the water pool was 120 cm and the height was 35 cm; a circular platform with a diameter of 4.5 cm and a height of 14.5 cm was placed at the center of the target quadrant (the third quadrant) (XR-XM101, Shanghai Xinruan Information Technology Co. Ltd, Shanghai, China). The MWM experiments were divided into initial spatial training and spatial probe test. All mice accepted four habituation trainings on day 0. The platform was visible (2 cm above the water surface), and the water was transparent. Before the initial spatial training test, the platform was hidden by submerging it 1.0 cm under the water surface (23-25°C), and the water was mixed with edible titanium dioxide pigments. On day 2, 4, and 6, the mice were carefully placed in the water maze and allowed to search for and climb onto the hidden platform for 60 s. If the mouse failed to find the platform within 60 s, it was guided to the platform, where it stayed for 30 s. Each mouse performed three trials per day, except the target quadrant, and the initial spatial training test lasted for three interval days. A video tracking system was used to record the escape latency to search for the hidden platform. The spatial probe test was implemented on the fourth day, with the hidden platform removed. Each mouse was released into different quadrants (except the target quadrant) and allowed to swim for 60 s. The data of the movement trajectories and number of platform crossings were recorded by a computerized video tracking system (Super Maze software, Shanghai Xinruan Information Technology Co., Ltd, China).

### Barnes maze test

The Barnes maze test for measuring spatial learning and memory was performed with appropriate modifications (Flores et al., 2018). Specifically, the maze platform (91 cm diameter, elevated 90 cm from the floor) consisted of 20 holes (each 5 cm in diameter). One target hole was equipped with an escape box; the other holes were not equipped with any escape devices. The mice were stimulated by strong light to find the escape box. During the adaptation period, each mouse explored the platform for 60 s. If the mouse found the escape box, it was allowed to stay there for another 60 s. Mice that did not find the escape box were guided to it and stayed there for 60 s. For the collection period, the goal was to train each mouse to find the target hole and enter the escape box within 180 s. Regardless of whether the mice found the escape box, they had to stay in the box for an additional 60 s. During this period, we recorded the head latency (primary latency) reaching the target hole. The protocol was conducted once a day for four consecutive days. In the probe test (day 5), each mouse was tested for 180 s. The target was in its original position, but the escape box was removed. The primary latency and the number of sniffing times (target preference) were recorded.

### Hematoxylin and eosin staining and Immunohistochemistry

Histopathology of colon slices was observed by H&E and IHC staining (Zhang *et al*., 2020). Colon sections were immersed in a 4% (*v*/*v*) paraformaldehyde for fixation and inserted in paraffin, then rehydrated with xylene and a gradient of ethanol (100%, 95%, 80%, and 75%). Then colon sections were washed in phosphate-buffered saline (PBS, pH 7.4) for 4 min. Tissue sections were stained with H&E and observed with an optical microscope. For IHC staining, sections were permeated in 0.5% Triton-X 100 for 15 min, and antigen retrieval was performed in citrate buffer. The sections were incubated with 3% H_2_O_2_ for 15 min to remove endogenous peroxidase. To prevent nonspecific staining, we used normal goat serum to block the sections for 2 h. The sections were incubated with the Claudin-1 primary antibody (1:300 dilution, Abcam Inc., Cambridge, MA, USA) at 4°C overnight. Then sections were incubated with the matching secondary horseradish peroxidase-streptavidin antibody (Streptavidin Peroxidase Link Detection Kits; Zhongshan Golden Bridge Biotechnology, Beijing, China). The sections were washed three times with PBS and then incubated with biotinylated goat anti-rabbit diluted in a secondary antibody dilution buffer. After staining nuclei by hematoxylin, neutral resin was used for sealing the sections. The stained tissue was observed with a light microscope (Olympus, Tokyo, Japan).

### Immunofluorescence staining

Deposits of Aβ and IBA-1 proten in the mouse brain were measured via immunofluorescence staining as described (Wang et al., 2019). Brain sections were deparaffinized and rehydrated, and then incubated with Aβ_1-42_ primary antibody (1:1600, Santa Cruz Inc., Dallas, Texas, USA) and IBA-1 primary antibody (1:1000 dilution, Abcam Inc., Cambridge, MA, USA) at 4°C overnight. After three washings with PBS, the sections were incubated with Alexa Fluor (555)-conjugated anti-rabbit secondary antibody for 20 min at 37°C. After washing six times with PBS, the sections were incubated with DAPI (4 μg/mL) for 30 min, washed, and covered. Immunofluorescence images were observed on an inverted fluorescence microscope (Olympus, Tokyo, Japan) (×200).

### Electron microscopy for structural analysis of postsynaptic density

Structural analysis of postsynaptic density was performed via electron microscopy (Zhao *et al*., 2019) of the CA1 region of the hippocampus. The hippocampus was split and treated in a cold fixative solution of 2.5% glutaraldehyde (pH 7.2) at 4°C for 4 h. After washing with PBS (0.1 mol/L, pH 7.2) three times, the specimens were post-fixed in 1% OsO_4_ (in 0.2 mol/L PBS, pH 7.2) at 4°C for 1 h and washed with PBS three times. The specimens were dehydrated for 15-20 min in a graded series of ethanol solutions (30, 50, 70, 80, 90, and 100%) and then transferred to acetone for 20 min. Materials were then permeated in an acetone-resin mixture (1:1) for 1 hour at 25°C and transferred to an acetone-resin mixture (1:3) overnight. Ultra-thin sections were placed in the regions which were close to the embedded blocks and kept away from the dorsal rim area, stained with uranyl acetate and alkaline lead citrate for 15 min, and then observed by transmission electron microscopy (JEOL, Tokyo, Japan) (×15000).

### Real-Time qPCR

Total RNA was isolated from brain tissues using Trizol reagent (Jingcai Bio., Xi’an, Shaanxi, China), and cDNA was generated using PrimeScript^TM^RT Master Mix reverse transcription kit (TaKaRa PrimeScript RT Master Mix, Dalian, China) and the CFX96^TM^ real-time system (Bio-Rad, Hercules, CA) (Zhang *et al*., 2020). All PCR analyses were performed in triplicate, and Ct values were quantified with corresponding standard curves. Relative quantitation of the target gene expression was calculated using the 2^-ΔΔCt^ method for statistical analysis. The primer sequences used are shown in Table S4.

### 16S rRNA Microbiome sequencing

Fecal samples were collected in a clean environment, and total cellular DNA was extracted with the E.Z.N.A. Stool DNA Kit (Omega, Norcross, GA, USA) according to the company instructions. The bacterial hypervariable V3-V4 region of 16S rRNA was chosen for MiSeq (Illumina, CA USA) paired-end 300 bp amplicon analysis using primers 341_F (5’-CCTACGGGNGGCWGCAG-3’) and 802_R (5’-TACNVGGGTATCTAATCC-3’). Library preparation followed a published method (Liu *et al*., 2020). The raw reads were merged and trimmed, chimeras were removed, and zero-radius Operational Taxonomic Units (zOTUs) with UNOISE was implemented in Search (v2.6.0). The green genes (13.8) 16s rRNA gene database was used as a reference for annotation. Detailed algorithms and parameters are given in Supplemental Materials.

A Kruskal-Wallis test was used to examine the influence of IF on the number of microbial species. Chao1 and ACE index reflected community richness, and the Shannon and Simpson index reflected community diversity. Principal coordinate analysis (PCoA) of the weighted and unweighted UniFrac distances or the Bray-Curtis dissimilarity was used to analyze the overall composition of gut microbiota. A permutational multivariate ANOVA (Adonis) (n=9999) and analysis of similarities were used to assess the effect of IF on PCoA scores of beta diversity metrics. To optimally identify a subset of microbial taxa affected by IF, we used a Partial Least Squares-Discrimination Analysis incorporated in the variable selection-within-repeated double cross-validation framework (VS-PLS-DA) on the auto-scaled relative abundance of microbiota to discriminate AD and WT mice groups vs. IF treated groups (i.e., AD+IF and WT+IF), AD mice group vs. AD+IF group, and WT mice group vs. WT+IF group, respectively (Shi et al., 2019). We further assessed IF-induced changes in the relative abundance of bacteria that were optimally selected by the VS-PLSDA modeling using ANOVA accounting for sex (females and males) and genotype (APP/PS1 and wild-type). FDR-p <0.05 was considered to be a significant difference. Rarified OTU data were used to predict functional genes with PICRUSt (v1.1.3). The predicted genes were annotated with KEGG at different levels, and the significantly abundant pathways were identified by edgeR with FDR-p <0.1. We also examined IF-induced changes in gut microbiota for male and female mice separately.

### Untargeted plasma metabolomics

After the animals were sacrificed, plasma samples were collected and stored at −80°C. The detection analysis was carried out by Shanghai Biotree Biomedical Technology Co., Ltd. Samples were analyzed using an ultra-performance liquid chromatography system (1290 UHPLC, Agilent) and a high-resolution tandem mass spectrometer Triple TOF 6600 (AB Sciex). Reverse-phase chromatography was employed, using both positive and negative electrospray ionization modes (ESI, RP+, and RP-). A 50 μL sample was mixed with 200 μL extract solution (acetonitrile: methanol = 1:1) containing internal standard (L-2-chlorophenylalanine, 2 μg/mL). After 30 s mixing, the samples were sonicated for 10 min in an ice-water bath, incubated at −40°C for 1 h, and centrifuged at 10000 rpm for 15 min at 4°C. A 220 μL sample of supernatant was transferred to a fresh tube and dried in a vacuum concentrator at 37°C. The dried samples were reconstituted in 200 μL of 50% acetonitrile by sonication on ice for 10 min, followed by centrifugation at 13000 rpm for 15 min at 4°C. A 75 μL sample of supernatant was transferred to a fresh glass vial for analysis. Samples were analyzed in one batch with a randomized injection order.

The ultra-performance liquid chromatography was conducted with a 1290 Infinity series UHPLC System (Agilent Technologies), equipped with a UPLC BEH Amide column (2.1*100 mm, 1.7 μm, Waters). The mobile phase consisted of 25 mmol/L ammonium acetate and 25 mmol/L ammonia hydroxide in water (pH 9.75) (A) and acetonitrile (B). The following elution gradient was used: 0∼0.5 min, 95% B; 0.5∼7.0 min, 95%∼65% B; 7.0∼8.0 min, 65%∼40% B; 8.0∼9.0 min, 40% B; 9.0∼9.1 min, 40%∼95% B; 9.1∼12.0 min, 95% B. The column temperature was 25°C. The auto-sampler temperature was 4°C, and the injection volume was 2 μL. The TripleTOF 6600 mass spectrometer (AB Sciex) was used to acquire MS/MS spectra on an information-dependent basis during an experiment. The acquisition software (Analyst TF 1.7, AB Sciex) continuously evaluated the full scan survey MS data as they were collected and triggered the acquisition of MS/MS spectra depending on preselected criteria. In each cycle, the 12 most intensive precursor ions with intensity above 100 were chosen for MS/MS at collision energy of 30 eV. The cycle time was 0.56 s. ESI source conditions were set as follows: Gas 1 as 60 psi, Gas 2 as 60 psi, Curtain Gas as 35 psi, Source Temperature as 600°C, Declustering potential as 60 V, Ion Spray Voltage Floating (ISVF) as 5000 V or −4000 V in positive or negative modes, respectively.

MS raw data (.wiff) files were converted to the mzXML format by ProteoWizard and processed by R package XCMS (version 3.2). The process included peak deconvolution, alignment, and integration. Minfrac and cut-off were set as 0.5 and 0.3, respectively. An in-house MS/MS database was used for metabolite identification. The stability and functionality of the system were monitored throughout all the instrumental analyses using quality controls, i.e. the pooling of all samples acquired at the beginning of the analytical sequence and after every 10 injections. We obtained 1936 and 1887 quantified metabolite features detected in RP+ and RP-, respectively, which were subjected to the VS-PLSDA modeling and ANOVA to investigate the effects of IF on plasma metabolome, as described above. The most relevant metabolite pathways induced by IF were revealed using the MetaboAnalyst (Pang et al., 2020).

### Integrated multi-omics analysis

Multivariate predictive modeling on each omics dataset was conducted using PLSDA incorporated into a repeated double cross-validations framework (rdCV-PLSDA). Outperforming the standard cross-validation, the double cross-validations procedure separates cross-validations into an outer “testing” loop and an inner “tuning” (or validation) loop to further reduce bias from overfitting models to experimental data. To gain a robust and reliable estimate of model performance, 200 repetitions of the outer cross-validations loop were performed, followed by permutation analysis (n=1000). A multivariate dimension reduction method, DIABLO (Data Integration Analysis for Biomarker discovery using a Latent component method for Omics), was employed for multiple omics integration (Rohart et al., 2017; Singh et al., 2019). Random use of a full design matrix was used to identify linear combinations of variables from each omics dataset that were maximally correlated. A tuning procedure was used to determine the optimal number of key variables in each dataset to be selected with a minimum misclassification rate. Model performance was then evaluated by 10-fold cross-validation.

### RNA Sequence Analysis

Brain tissue from the hippocampus was macro-dissected and stored at −80°C. Total RNA was extracted using TriZol (Invitrogen, Carlsbad, UA, USA) according to the manufacturer’s instructions, followed by being qualified and quantified using a NanoDrop and Agilent 2100 bioanalyzer (Thermo Fisher Scientific, MA, USA). RNA sequencing libraries were prepared using BGISEQ-500 (BGI-Shenzhen, China). Sequencing data were filtered and trimmed using Trimmomatic v0.38 (Bolger, Lohse, & Usadel, 2014) to obtain high-quality clean-read data. Clean reads were mapped to the *Mus musculus* genome sequence (ftp://ftp.ncbi.nlm.nih.gov/genomes/all/GCF/000/001/635/GCF_000001635.26_GRCm38.p6) using Hisat2 v2-2.1.0 (Kim, Langmead, & Salzberg, 2015). The reads of each sample were then assembled into transcripts and compared with reference gene models using StringTie v1.3.4d (M. Pertea et al., 2015). We merged the 55 transcripts to obtain a consensus transcript using a StringTie-Merge program. Transcripts that did not exist in the CDS database of the *Mus musculus* genome were extracted to predict new genes. The gene expression FPKM values were calculated using the StringTie Merge program based on the consensus transcript. We then used VS-PLSDA modeling and ANOVA to identify significant IF-induced changes in gene expression. KEGG and Gene Ontology enrichment analyses were performed using the WebGestalt 2019 (WEB-based Gene Set Analysis Toolkit) (Liao et al., 2019). The most relevant genes associated with IF were further filtered according to KEGG about Alzheimer’s disease, oxidative phosphorylation, neurotrophin signaling pathway, and synapse (glutamatergic, cholinergic, serotonergic, GABAergic, and dopaminergic). The expression of these genes was subjected to the multi-omics analysis integrating with the relative abundance of gut microbiota and plasma levels of metabolites that were affected by IF treatment.

### Statistical analysis

Data were reported as mean ± SEM of at least three independent experiments and analyzed with Newman-Keuls test for multiple comparison test by Graphpad Prism 6.0 software (GraphPad Software Inc., San Diego, CA). Significant differences between mean values were determined by Two-way ANOVA with genotype (wild-type and APP/PS1) and IF (*ad libitum* diet /IF diet) as factors. Statistical significance was presented as mean ± SEM. *p < 0.05, **p < 0.01, compared with WT group, ^#^p < 0.05, ^##^p < 0.01 compared with the AD group.

