## Supplementary figures for "Intermittent fasting mitigates cognitive deficits in Alzheimer’s disease via the gut-brain axis"

### Supplementary Figure Legends

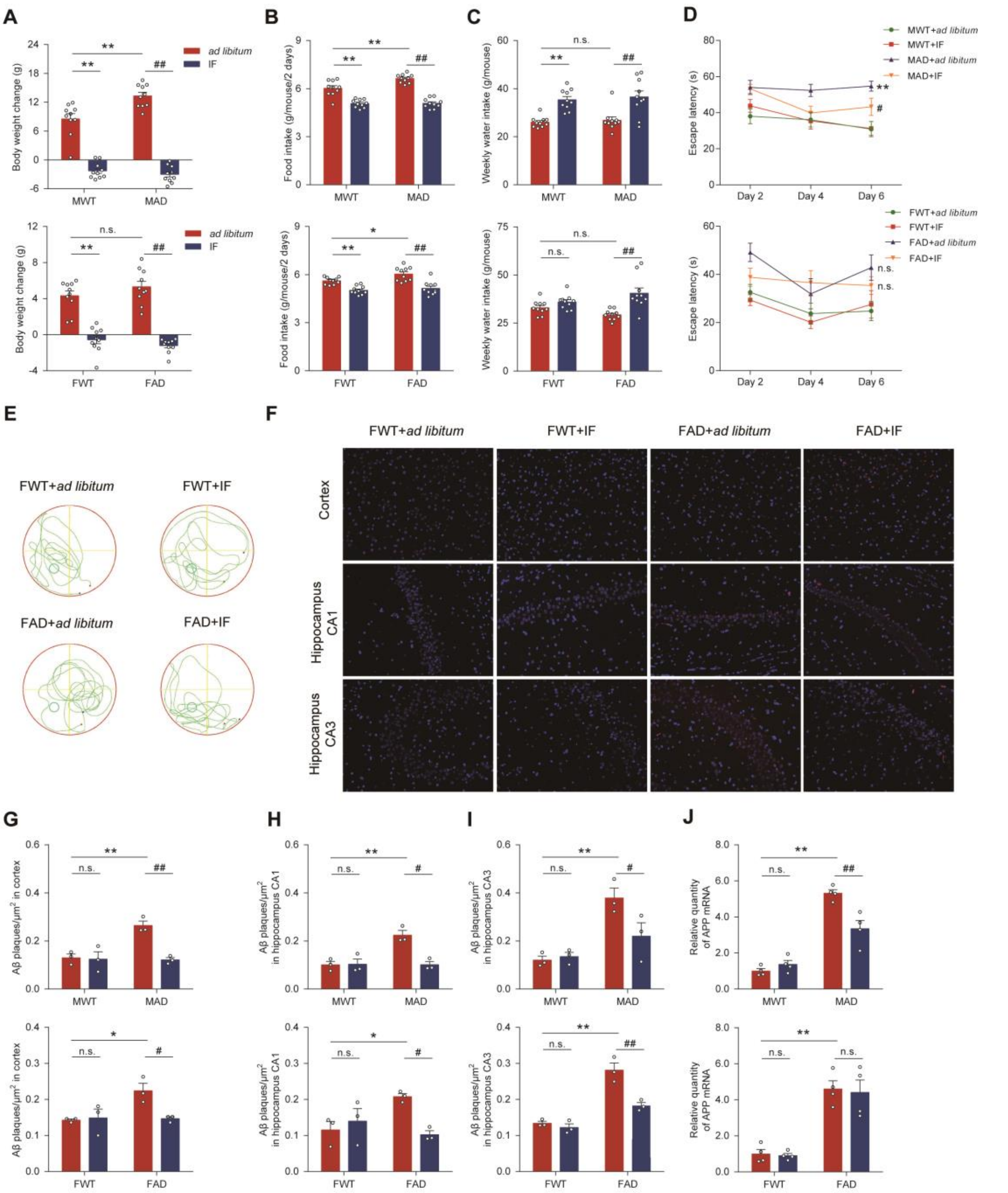

**Supplementary Fig. 1. IF ameliorated cognitive impairments behavior in the AD mice.**

**(A)** Body weight change of males and females. **(B)** Food intake of males and females. **(C)** Weekly water intake of males and females. **(D)** Escape latency of males and females. (**A-D**, n = 10 mice per group). **(E)** Mouse trajectory of female mice. **(F)** Representative images of Aβ plaques in cortex, CA1 and CA3 region of the hippocampus of female mice; Scale bars: 100 μm.

**(G-I)** Aβ plaque density (number of plaques/$\mu m^2$) in the cortex, CA1, and CA3 region of the hippocampus between males and females (n=3 mice per group). **(J)** The mRNA levels of *APP* in the hippocampus of males and females. (n=4 mice per group). Data presented as mean ± SEM. $^{*}p < 0.05$, $^{**}p < 0.01$, compared with the male/female WT+*ad libitum* group, $^{\#}p < 0.05$, $^{\#\#}p < 0.01$ compared with the male/female AD+*ad libitum* group. Significant differences between mean values were determined by two-way ANOVA with Newman-Keuls multiple comparisons test.

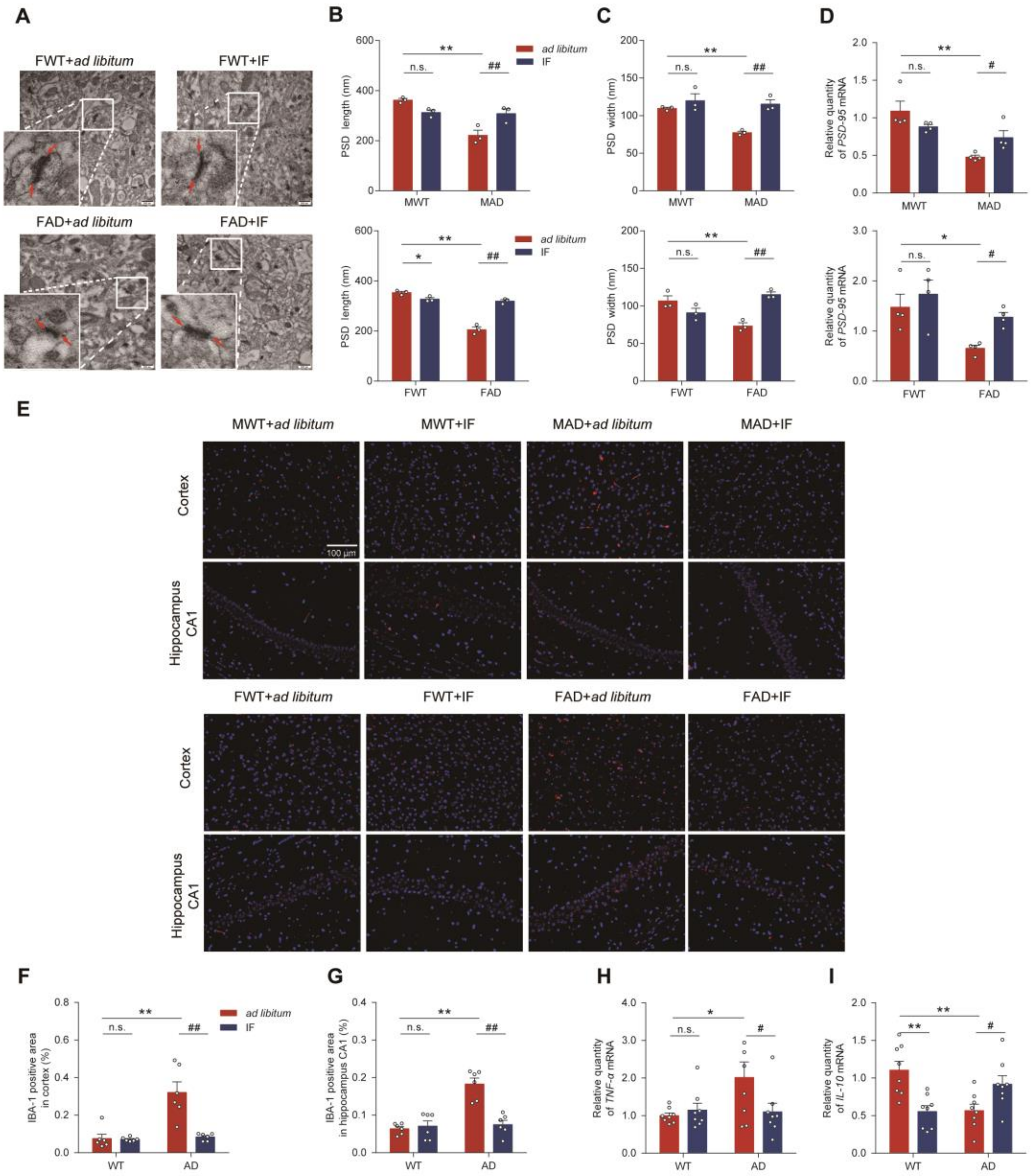


**Supplementary Fig. 2. IF improved synaptic ultrastructure and neuroinflammation in AD mice**.

**(A)** Representative images of the ultrastructure of synapse in the hippocampus of female mice. **(B-C)** The length and width of PSD in the hippocampus of males and females (n=3 mice per group). **(D)** The mRNA levels of *PSD-95* in the hippocampus of males and females. (n=4 mice per group). **(E)** Representative images of immunohistochemical staining of IBA-1 protein expression in the cortex and CA1 region of the hippocampus; representative images of immunohistochemical staining were captured from 3 slices from 3 mice per group. Scale bars: 100 μm. **(F-G)** Quantitative analysis of IBA-1 immunofluorescence staining. **(H-I)** The mRNA levels of *TNF-α* and *IL-10* in the hippocampus (n=8, male and female in half). Data presented as mean ± SEM. $^{*}p < 0.05$, $^{**}p < 0.01$, compared with the male/female WT+*ad libitum* group, $^{\#}p < 0.05$, $^{\#\#}p < 0.01$ compared with the male/female AD+*ad libitum* group. Significant differences between mean values were determined by two-way ANOVA with Newman-Keuls multiple comparisons test.

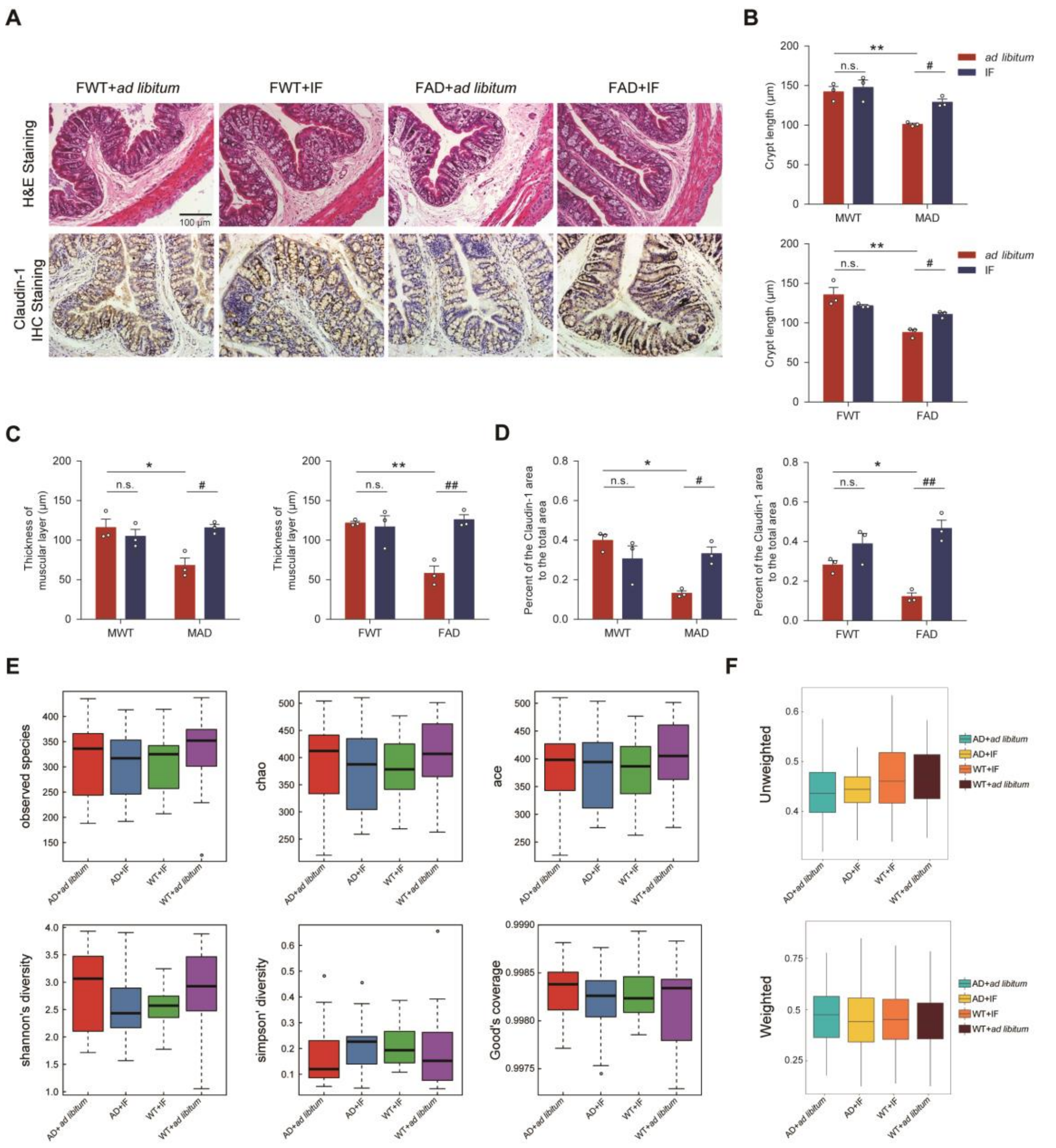

**Supplementary Fig. 3. IF improved the gut barrier and restructured the gut microbiota of AD mice.**

**(A)** Representative images of H&E staining and Claudin-1 protein IHC of the colon of female mice; Scale bars: 100 μm. **(B-D)** The crypt length, the thickness of the muscular layer, and the proportion of the Claudin-1 area in the total area in males and females (n = 3 mice per group). **(E-F)** Alpha diversity box and Beta diversity index box (n = 18 mice per group). The five lines from bottom to top are minimum, first quartile, median, third quartile, and maximum.

Data presented as mean ± SEM. $^{*}p < 0.05$, $^{**}p < 0.01$, compared with the male/female WT+*ad libitum* group, $^{\#}p < 0.05$, $^{\#\#}p < 0.01$ compared with the male/female AD+*ad libitum* group. Significant differences between mean values were determined by two-way ANOVA with Newman-Keuls multiple comparisons test.

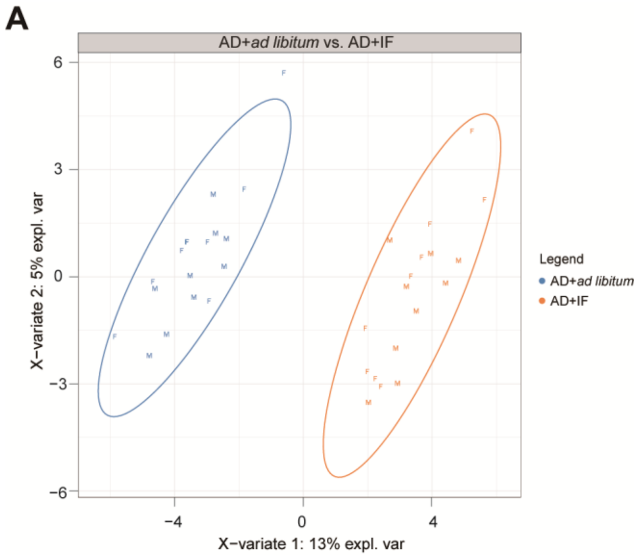


**Supplementary Fig. 4. A score plot revealing a clear discrimination between AD+*ad libitum* and AD+IF group.**

**(A)** The plot was obtained by using partial least squares regression-discrimination analysis on a selection of IF- related 6 genera and 28 OTUs that were individually differential between AD+*ad libitum* and AD+IF group.

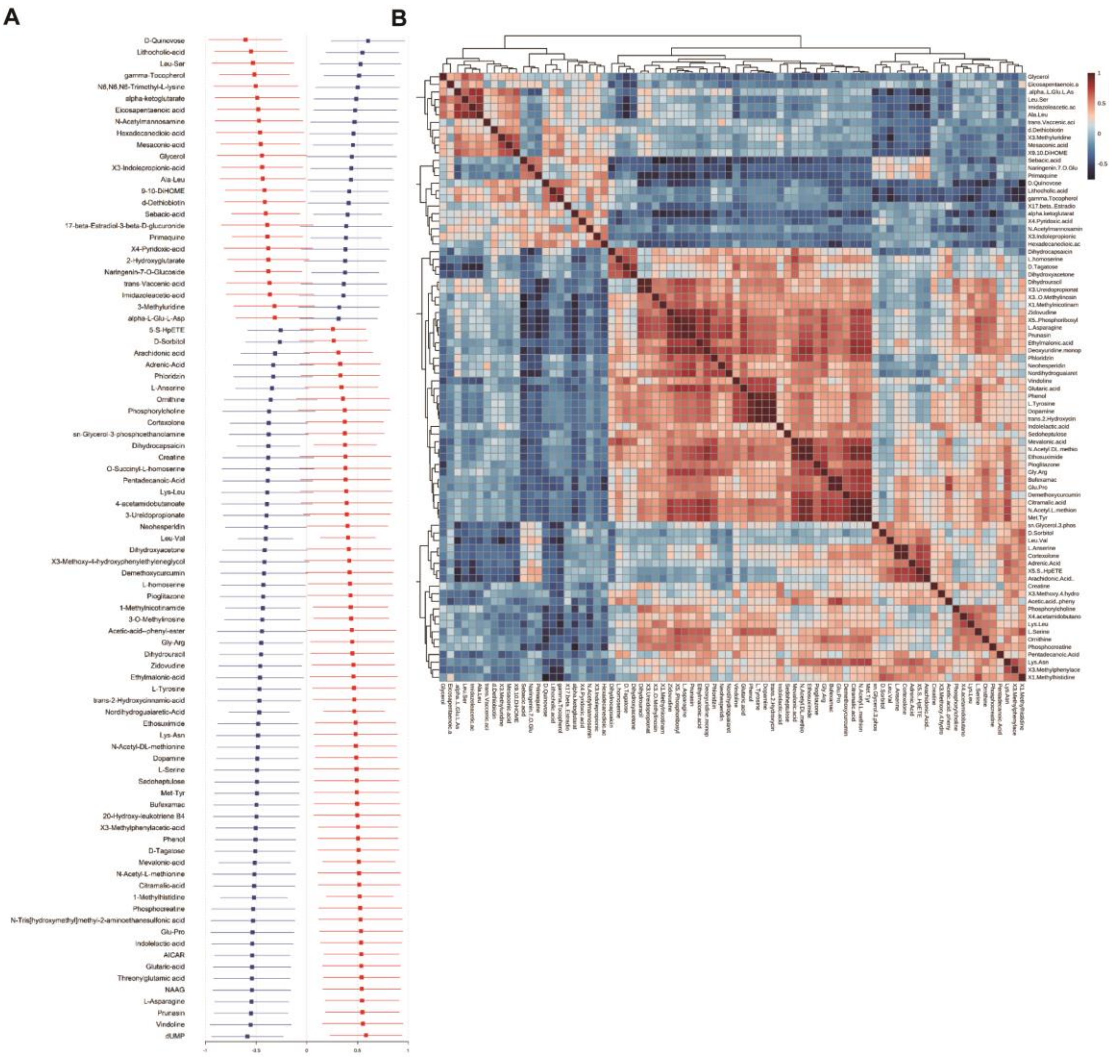


**Supplementary Fig. 5. IF induced changes in plasma metabolites.**

**(A)** Differences in a selection of 87 metabolites between AD+*ad libitum* and AD+IF group, as assessed by ANOVA. Least square means ± 95% confident intervals are presented. **(B)** A z-scored heatmap showing inter-correlations between 87 metabolites.

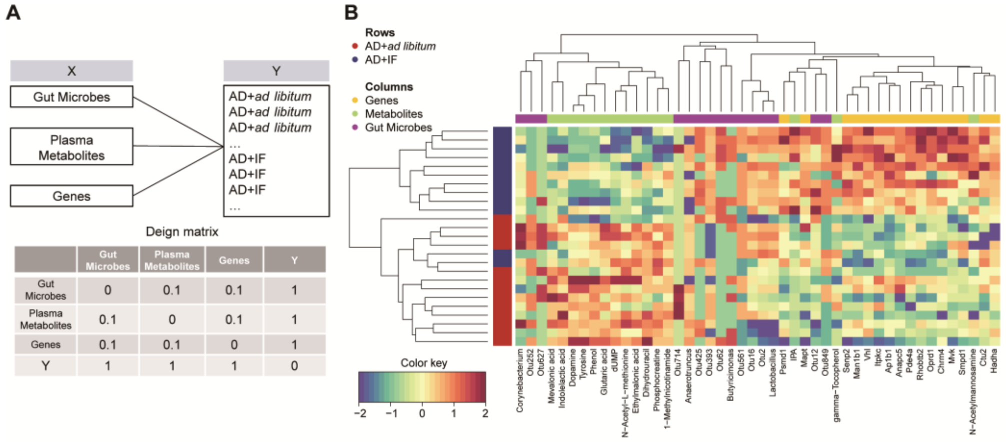


**Supplementary Fig. 6. DIABLO model for integrating 175 differential AD+*ad libitum* vs. AD+IF genes expressed in the hippocampus, 34 gut bacteria and 87 plasma metabolites.**

**(A)** Matrix design for the DIABLO analysis. **(B)** A heatmap showing the clusters complete linkage of the multi-OMICs signature that were demonstrated by using Euclidean distance. Samples are represented in rows, selected features on the first component in columns.

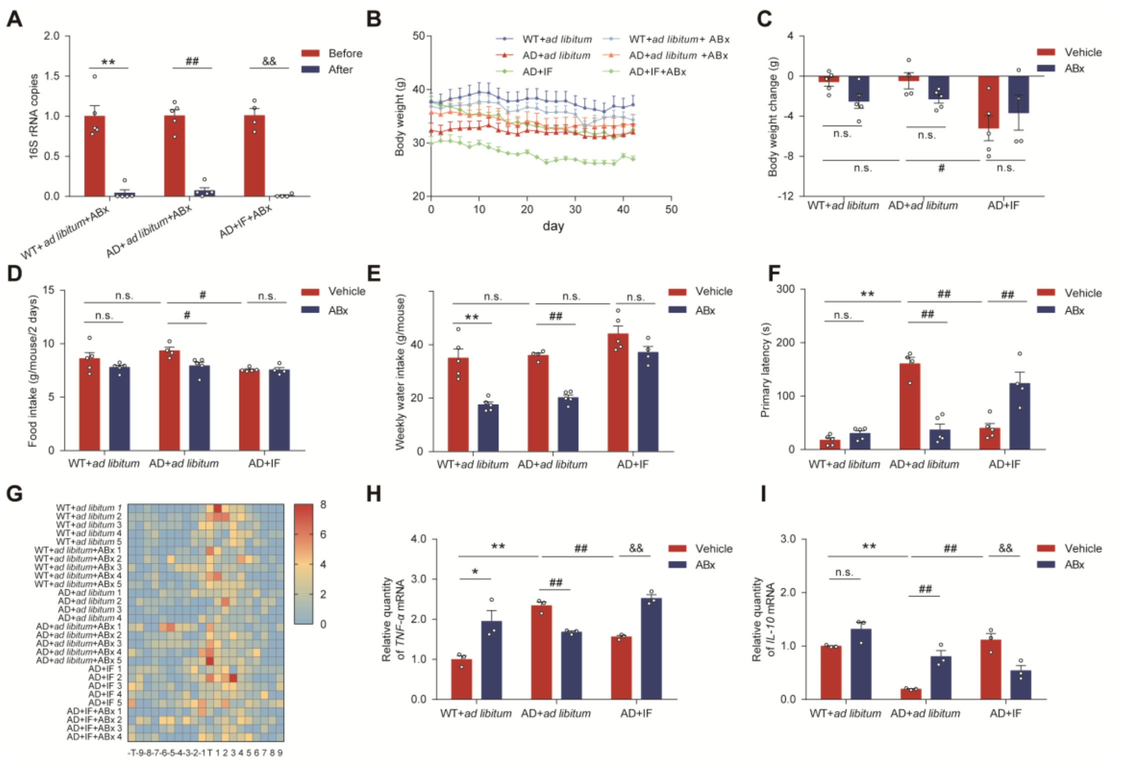


**Supplementary Fig. 7. The improvement of cognitive function in the AD mice by IF was mediated by intestinal microbiota.**

The mice were administrated with antibiotics in the drinking water starting 14 days before the 4-week IF regimen and throughout the experiment (the detailed antibiotics treatment is as described in “Methods”). **(A)** 16S rRNA copies (WT+*ad libitum*+ABx, AD+*ad libitum*+ABx: n=5; AD+IF+ABx: n=4). **(B)** and **(C)** Body weight changes. **(D)** Food intake. **(E)** Weekly water intake. **(F)** Primary latency in probe trial of Barnes maze. **(G)** Quantitative assessment of the number of times each mouse poked its head into each hole of the Barnes maze (WT+*ad libitum*, WT+*ad libitum*+ABx, AD+*ad libitum*+ABx, AD+IF: n=5; AD+*ad libitum*, AD+IF+ABx: n=4). **(H)** The mRNA levels of *TNF-α* in the hippocampus (n=3). **(I)** The mRNA levels of *IL-10* in the hippocampus (n=3). Data presented as mean ± SEM. $^{*}p < 0.05$, $^{**}p < 0.01$, compared with the male/female WT+*ad libitum* group, $^{\#}p < 0.05$, $^{\#\#}p < 0.01$ compared with the male/female AD+*ad libitum* group, $^{\&}p < 0.05$, $^{\&\&}p < 0.01$ compared with the AD+IF group. Significant differences between mean values were determined by two-way ANOVA with Newman-Keuls multiple comparisons test.
