## Supplementary tables for "Intermittent fasting mitigates cognitive deficits in Alzheimer’s disease via the gut-brain axis"

### **Supplementary Table 1.** 280 KEGG pathways

| | geneSet | description | size | overlap | expect | enrichment Ratio | pValue | FDR |
|---|---|---|---|---|---|---|---|---|
| IF-upregulated | mmu04310 | Wnt signaling pathway | 148 | 16 | 4.578610 | 3.494510 | 0.000012 | 0.00203 |
| IF-upregulated | mmu05224 | Breast cancer | 147 | 16 | 4.547674 | 3.518282 | 0.000011 | 0.00203 |
| IF-upregulated | mmu05225 | Hepatocellular carcinoma | 171 | 16 | 5.290151 | 3.024488 | 0.000075 | 0.00810 |
| IF-upregulated | mmu04150 | mTOR signaling pathway | 153 | 13 | 4.733293 | 2.746502 | 0.000888 | 0.04349 |
| IF-upregulated | mmu04550 | Signaling pathways regulating pluripotency of stem cells | 137 | 12 | 4.238308 | 2.831318 | 0.001067 | 0.04349 |
| IF-upregulated | mmu05217 | Basal cell carcinoma | 63 | 8 | 1.949003 | 4.104663 | 0.000667 | 0.04349 |
| IF-upregulated | mmu05226 | Gastric cancer | 150 | 13 | 4.640483 | 2.801432 | 0.000737 | 0.04349 |
| IF-upregulated | mmu04916 | Melanogenesis | 100 | 10 | 3.093656 | 3.232422 | 0.001008 | 0.04349 |
| IF-upregulated | mmu05213 | Endometrial cancer | 58 | 7 | 1.794320 | 3.901199 | 0.001952 | 0.07071 |
| IF-upregulated | mmu04140 | Autophagy | 130 | 11 | 4.021752 | 2.735126 | 0.002255 | 0.07351 |
| IF-upregulated | mmu04390 | Hippo signaling pathway | 154 | 12 | 4.764230 | 2.518770 | 0.002892 | 0.08571 |
| IF-upregulated | mmu04330 | Notch signaling pathway | 49 | 6 | 1.515891 | 3.958068 | 0.003788 | 0.10291 |
| IF-upregulated | mmu04010 | MAPK signaling pathway | 294 | 18 | 9.095347 | 1.979034 | 0.004378 | 0.10333 |
| IF-upregulated | mmu04071 | Sphingolipid signaling pathway | 122 | 10 | 3.774260 | 2.649526 | 0.004437 | 0.10333 |
| IF-upregulated | mmu04917 | Prolactin signaling pathway | 72 | 7 | 2.227432 | 3.142632 | 0.006637 | 0.11388 |
| IF-upregulated | mmu05216 | Thyroid cancer | 37 | 5 | 1.144653 | 4.368138 | 0.005284 | 0.11388 |
| IF-upregulated | mmu05210 | Colorectal cancer | 88 | 8 | 2.722417 | 2.938565 | 0.005713 | 0.11388 |
| IF-upregulated | mmu05214 | Glioma | 71 | 7 | 2.196495 | 3.186895 | 0.006150 | 0.11388 |
| IF-upregulated | mmu05218 | Melanoma | 72 | 7 | 2.227432 | 3.142632 | 0.006637 | 0.11388 |
| IF-upregulated | mmu04919 | Thyroid hormone signaling pathway | 115 | 9 | 3.557704 | 2.529721 | 0.009146 | 0.14907 |
| IF-upregulated | mmu05215 | Prostate cancer | 97 | 8 | 3.000846 | 2.665915 | 0.010138 | 0.15738 |
| IF-upregulated | mmu05231 | Choline metabolism in cancer | 99 | 8 | 3.062719 | 2.612058 | 0.011396 | 0.16886 |
| IF-upregulated | mmu05200 | Pathways in cancer | 530 | 26 | 16.396375 | 1.585716 | 0.012807 | 0.18153 |
| IF-upregulated | mmu03013 | RNA transport | 167 | 11 | 5.166405 | 2.129140 | 0.014426 | 0.19595 |
| IF-upregulated | mmu05223 | Non-small cell lung cancer | 66 | 6 | 2.041813 | 2.938565 | 0.015946 | 0.20793 |
| IF-upregulated | mmu04360 | Axon guidance | 175 | 11 | 5.413897 | 2.031808 | 0.019786 | 0.24809 |
| IF-upregulated | mmu04211 | Longevity regulating pathway | 90 | 7 | 2.784290 | 2.514106 | 0.021126 | 0.25507 |
| IF-upregulated | mmu01522 | Endocrine resistance | 93 | 7 | 2.877100 | 2.433006 | 0.024797 | 0.28871 |
| IF-upregulated | mmu04015 | Rap1 signaling pathway | 209 | 12 | 6.465740 | 1.855936 | 0.028731 | 0.32297 |
| IF-upregulated | mmu00900 | Terpenoid backbone biosynthesis | 23 | 3 | 0.711541 | 4.216202 | 0.032815 | 0.34248 |
| IF-upregulated | mmu00260 | Glycine, serine and threonine metabolism | 40 | 4 | 1.237462 | 3.232422 | 0.034197 | 0.34248 |
| IF-upregulated | mmu04370 | VEGF signaling pathway | 58 | 5 | 1.794320 | 2.786571 | 0.032985 | 0.34248 |
| IF-upregulated | mmu04810 | Regulation of actin cytoskeleton | 215 | 12 | 6.651360 | 1.804142 | 0.034668 | 0.34248 |
| IF-upregulated | mmu01521 | EGFR tyrosine kinase inhibitor resistance | 80 | 6 | 2.474924 | 2.424316 | 0.037025 | 0.35500 |
| IF-upregulated | mmu04066 | HIF-1 signaling pathway | 105 | 7 | 3.248338 | 2.154948 | 0.043810 | 0.40806 |
| IF-upregulated | mmu04928 | Parathyroid hormone synthesis, secretion and action | 107 | 7 | 3.310211 | 2.114669 | 0.047695 | 0.43190 |
| IF-upregulated | mmu04068 | FoxO signaling pathway | 132 | 8 | 4.083625 | 1.959044 | 0.051927 | 0.44286 |
| IF-upregulated | mmu03050 | Proteasome | 46 | 4 | 1.423082 | 2.810802 | 0.052981 | 0.44286 |
| IF-upregulated | mmu05205 | Proteoglycans in cancer | 204 | 11 | 6.311057 | 1.742973 | 0.051689 | 0.44286 |
| IF-upregulated | mmu05211 | Renal cell carcinoma | 68 | 5 | 2.103686 | 2.376781 | 0.058749 | 0.47483 |
| IF-upregulated | mmu04914 | Progesterone-mediated oocyte maturation | 90 | 6 | 2.784290 | 2.154948 | 0.059718 | 0.47483 |
| IF-upregulated | mmu05221 | Acute myeloid leukemia | 69 | 5 | 2.134622 | 2.342335 | 0.061816 | 0.47981 |
| IF-upregulated | mmu00520 | Amino sugar and nucleotide sugar metabolism | 49 | 4 | 1.515891 | 2.638712 | 0.064102 | 0.48598 |
| IF-upregulated | mmu04120 | Ubiquitin mediated proteolysis | 139 | 8 | 4.300181 | 1.860387 | 0.066288 | 0.49113 |
| IF-upregulated | mmu04115 | p53 signaling pathway | 71 | 5 | 2.196495 | 2.276353 | 0.068216 | 0.49419 |
| IF-upregulated | mmu04520 | Adherens junction | 72 | 5 | 2.227432 | 2.244737 | 0.071550 | 0.50707 |
| IF-upregulated | mmu01212 | Fatty acid metabolism | 52 | 4 | 1.608701 | 2.486478 | 0.076346 | 0.52955 |
| IF-upregulated | mmu04070 | Phosphatidylinositol signaling system | 98 | 6 | 3.031782 | 1.979034 | 0.082752 | 0.54931 |

| | | | | | | | | |
|---|---|---|---|---|---|---|---|---|
| IF-upregulated | mmu04722 | Neurotrophin signaling pathway | 121 | 7 | 3.743323 | 1.869996 | 0.080961 | 0.54931 |
| IF-upregulated | mmu04620 | Toll-like receptor signaling pathway | 99 | 6 | 3.062719 | 1.959044 | 0.085935 | 0.54931 |
| IF-upregulated | mmu05220 | Chronic myeloid leukemia | 76 | 5 | 2.351178 | 2.126593 | 0.085757 | 0.54931 |
| IF-upregulated | mmu00051 | Fructose and mannose metabolism | 35 | 3 | 1.082779 | 2.770647 | 0.092842 | 0.55689 |
| IF-upregulated | mmu00280 | Valine, leucine and isoleucine degradation | 56 | 4 | 1.732447 | 2.308873 | 0.094352 | 0.55689 |
| IF-upregulated | mmu00450 | Selenocompound metabolism | 17 | 2 | 0.525921 | 3.802849 | 0.095623 | 0.55689 |
| IF-upregulated | mmu04152 | AMPK signaling pathway | 126 | 7 | 3.898006 | 1.795790 | 0.095441 | 0.55689 |
| IF-upregulated | mmu05206 | MicroRNAs in cancer | 281 | 13 | 8.693172 | 1.495426 | 0.095938 | 0.55689 |
| IF-upregulated | mmu05166 | Human T-cell leukemia virus 1 infection | 282 | 13 | 8.724109 | 1.490124 | 0.097934 | 0.55689 |
| IF-upregulated | mmu05143 | African trypanosomiasis | 36 | 3 | 1.113716 | 2.693685 | 0.099079 | 0.55689 |
| IF-upregulated | mmu04014 | Ras signaling pathway | 233 | 11 | 7.208218 | 1.526036 | 0.107461 | 0.57753 |
| IF-upregulated | mmu00511 | Other glycan degradation | 18 | 2 | 0.556858 | 3.591580 | 0.105460 | 0.57753 |
| IF-upregulated | mmu00310 | Lysine degradation | 59 | 4 | 1.825257 | 2.191472 | 0.109051 | 0.57753 |
| IF-upregulated | mmu05165 | Human papillomavirus infection | 370 | 16 | 11.446526 | 1.397804 | 0.109837 | 0.57753 |
| IF-upregulated | mmu00620 | Pyruvate metabolism | 38 | 3 | 1.175589 | 2.551912 | 0.112037 | 0.57975 |
| IF-upregulated | mmu00290 | Valine, leucine and isoleucine biosynthesis | 4 | 1 | 0.123746 | 8.081055 | 0.118142 | 0.58090 |
| IF-upregulated | mmu00561 | Glycerolipid metabolism | 61 | 4 | 1.887130 | 2.119621 | 0.119387 | 0.58090 |
| IF-upregulated | mmu04934 | Cushing syndrome | 158 | 8 | 4.887976 | 1.636669 | 0.116558 | 0.58090 |
| IF-upregulated | mmu04012 | ErbB signaling pathway | 84 | 5 | 2.598671 | 1.924061 | 0.118219 | 0.58090 |
| IF-upregulated | mmu04213 | Longevity regulating pathway | 62 | 4 | 1.918066 | 2.085433 | 0.124708 | 0.59787 |
| IF-upregulated | mmu04141 | Protein processing in endoplasmic reticulum | 163 | 8 | 5.042659 | 1.586465 | 0.132474 | 0.59961 |
| IF-upregulated | mmu04725 | Cholinergic synapse | 113 | 6 | 3.495831 | 1.716330 | 0.137295 | 0.59961 |
| IF-upregulated | mmu04910 | Insulin signaling pathway | 140 | 7 | 4.331118 | 1.616211 | 0.142998 | 0.59961 |
| IF-upregulated | mmu04151 | PI3K-Akt signaling pathway | 358 | 15 | 11.075287 | 1.354367 | 0.143186 | 0.59961 |
| IF-upregulated | mmu04912 | GnRH signaling pathway | 90 | 5 | 2.784290 | 1.795790 | 0.145840 | 0.59961 |
| IF-upregulated | mmu04540 | Gap junction | 86 | 5 | 2.660544 | 1.879315 | 0.127131 | 0.59961 |
| IF-upregulated | mmu04973 | Carbohydrate digestion and absorption | 43 | 3 | 1.330272 | 2.255178 | 0.146979 | 0.59961 |
| IF-upregulated | mmu03430 | Mismatch repair | 22 | 2 | 0.680604 | 2.938565 | 0.147145 | 0.59961 |
| IF-upregulated | mmu00860 | Porphyrin and chlorophyll metabolism | 41 | 3 | 1.268399 | 2.365187 | 0.132597 | 0.59961 |
| IF-upregulated | mmu05219 | Bladder cancer | 41 | 3 | 1.268399 | 2.365187 | 0.132597 | 0.59961 |
| IF-upregulated | mmu05230 | Central carbon metabolism in cancer | 64 | 4 | 1.979940 | 2.020264 | 0.135645 | 0.59961 |
| IF-upregulated | mmu04670 | Leukocyte transendothelial migration | 115 | 6 | 3.557704 | 1.686481 | 0.145609 | 0.59961 |
| IF-upregulated | mmu05140 | Leishmaniasis | 67 | 4 | 2.072749 | 1.929804 | 0.152747 | 0.61404 |
| IF-upregulated | mmu05161 | Hepatitis B | 143 | 7 | 4.423927 | 1.582304 | 0.154453 | 0.61404 |
| IF-upregulated | mmu04664 | Fc epsilon RI signaling pathway | 68 | 4 | 2.103686 | 1.901425 | 0.158621 | 0.62302 |
| IF-upregulated | mmu00592 | alpha-Linolenic acid metabolism | 25 | 2 | 0.773414 | 2.585938 | 0.180225 | 0.64816 |
| IF-upregulated | mmu00340 | Histidine metabolism | 24 | 2 | 0.742477 | 2.693685 | 0.169066 | 0.64816 |
| IF-upregulated | mmu04072 | Phospholipase D signaling pathway | 147 | 7 | 4.547674 | 1.539249 | 0.170363 | 0.64816 |
| IF-upregulated | mmu04110 | Cell cycle | 123 | 6 | 3.805196 | 1.576791 | 0.181031 | 0.64816 |
| IF-upregulated | mmu00564 | Glycerophospholipid metabolism | 97 | 5 | 3.000846 | 1.666197 | 0.181175 | 0.64816 |
| IF-upregulated | mmu04714 | Thermogenesis | 230 | 10 | 7.115408 | 1.405401 | 0.175417 | 0.64816 |
| IF-upregulated | mmu05169 | Epstein-Barr virus infection | 230 | 10 | 7.115408 | 1.405401 | 0.175417 | 0.64816 |
| IF-upregulated | mmu04662 | B cell receptor signaling pathway | 72 | 4 | 2.227432 | 1.795790 | 0.182916 | 0.64816 |
| IF-upregulated | mmu05203 | Viral carcinogenesis | 231 | 10 | 7.146344 | 1.399317 | 0.178704 | 0.64816 |
| IF-upregulated | mmu04142 | Lysosome | 124 | 6 | 3.836133 | 1.564075 | 0.185686 | 0.65090 |
| IF-upregulated | mmu00510 | N-Glycan biosynthesis | 49 | 3 | 1.515891 | 1.979034 | 0.192801 | 0.66865 |
| IF-upregulated | mmu00071 | Fatty acid degradation | 50 | 3 | 1.546828 | 1.939453 | 0.200763 | 0.67477 |
| IF-upregulated | mmu00650 | Butanoate metabolism | 27 | 2 | 0.835287 | 2.394387 | 0.202846 | 0.67477 |

| | | | | | | | | |
|---|---|---|---|---|---|---|---|---|
| IF-upregulated | mmu04933 | AGE-RAGE signaling pathway in diabetic complications | 100 | 5 | 3.093656 | 1.616211 | 0.197216 | 0.67477 |
| IF-upregulated | mmu00601 | Glycosphingolipid biosynthesis | 27 | 2 | 0.835287 | 2.394387 | 0.202846 | 0.67477 |
| IF-upregulated | mmu05202 | Transcriptional misregulation in cancer | 183 | 8 | 5.661390 | 1.413081 | 0.206131 | 0.67877 |
| IF-upregulated | mmu03060 | Protein export | 28 | 2 | 0.866224 | 2.308873 | 0.214274 | 0.68391 |
| IF-upregulated | mmu04122 | Sulfur relay system | 8 | 1 | 0.247492 | 4.040527 | 0.222374 | 0.68391 |
| IF-upregulated | mmu04218 | Cellular senescence | 186 | 8 | 5.754199 | 1.390289 | 0.218391 | 0.68391 |
| IF-upregulated | mmu04064 | NF-kappa B signaling pathway | 104 | 5 | 3.217402 | 1.554049 | 0.219327 | 0.68391 |
| IF-upregulated | mmu04926 | Relaxin signaling pathway | 131 | 6 | 4.052689 | 1.480499 | 0.219516 | 0.68391 |
| IF-upregulated | mmu05132 | Salmonella infection | 78 | 4 | 2.413051 | 1.657652 | 0.221417 | 0.68391 |
| IF-upregulated | mmu00740 | Riboflavin metabolism | 8 | 1 | 0.247492 | 4.040527 | 0.222374 | 0.68391 |
| IF-upregulated | mmu05160 | Hepatitis C | 134 | 6 | 4.145498 | 1.447353 | 0.234616 | 0.71481 |
| IF-upregulated | mmu01523 | Antifolate resistance | 30 | 2 | 0.928097 | 2.154948 | 0.237293 | 0.71627 |
| IF-upregulated | mmu04923 | Regulation of lipolysis in adipocytes | 55 | 3 | 1.701511 | 1.763139 | 0.241607 | 0.72260 |
| IF-upregulated | mmu04210 | Apoptosis | 136 | 6 | 4.207372 | 1.426068 | 0.244858 | 0.72567 |
| IF-upregulated | mmu00640 | Propanoate metabolism | 31 | 2 | 0.959033 | 2.085433 | 0.248854 | 0.73087 |
| IF-upregulated | mmu00410 | beta-Alanine metabolism | 32 | 2 | 0.989970 | 2.020264 | 0.260432 | 0.74474 |
| IF-upregulated | mmu04136 | Autophagy | 32 | 2 | 0.989970 | 2.020264 | 0.260432 | 0.74474 |
| IF-upregulated | mmu00062 | Fatty acid elongation | 32 | 2 | 0.989970 | 2.020264 | 0.260432 | 0.74474 |
| IF-upregulated | mmu04510 | Focal adhesion | 199 | 8 | 6.156375 | 1.299466 | 0.274385 | 0.77782 |
| IF-upregulated | mmu04666 | Fc gamma R-mediated phagocytosis | 87 | 4 | 2.691480 | 1.486171 | 0.282530 | 0.78992 |
| IF-upregulated | mmu04114 | Oocyte meiosis | 115 | 5 | 3.557704 | 1.405401 | 0.283499 | 0.78992 |
| IF-upregulated | mmu04730 | Long-term depression | 61 | 3 | 1.887130 | 1.589716 | 0.292153 | 0.80543 |
| IF-upregulated | mmu03030 | DNA replication | 35 | 2 | 1.082779 | 1.847098 | 0.295154 | 0.80543 |
| IF-upregulated | mmu00590 | Arachidonic acid metabolism | 89 | 4 | 2.753353 | 1.452774 | 0.296477 | 0.80543 |
| IF-upregulated | mmu04614 | Renin-angiotensin system | 36 | 2 | 1.113716 | 1.795790 | 0.306688 | 0.82297 |
| IF-upregulated | mmu04137 | Mitophagy | 63 | 3 | 1.949003 | 1.539249 | 0.309194 | 0.82297 |
| IF-upregulated | mmu04657 | IL-17 signaling pathway | 91 | 4 | 2.815227 | 1.420845 | 0.310508 | 0.82297 |
| IF-upregulated | mmu05222 | Small cell lung cancer | 92 | 4 | 2.846163 | 1.405401 | 0.317549 | 0.83352 |
| IF-upregulated | mmu04723 | Retrograde endocannabinoid signaling | 150 | 6 | 4.640483 | 1.292969 | 0.319601 | 0.83352 |
| IF-upregulated | mmu04960 | Aldosterone-regulated sodium reabsorption | 38 | 2 | 1.175589 | 1.701275 | 0.329638 | 0.85287 |
| IF-upregulated | mmu03450 | Non-homologous end-joining | 13 | 1 | 0.402175 | 2.486478 | 0.335572 | 0.86139 |
| IF-upregulated | mmu04611 | Platelet activation | 124 | 5 | 3.836133 | 1.303396 | 0.338368 | 0.86178 |
| IF-upregulated | mmu04720 | Long-term potentiation | 67 | 3 | 2.072749 | 1.447353 | 0.343342 | 0.86398 |
| IF-upregulated | mmu04750 | Inflammatory mediator regulation of TRP channels | 125 | 5 | 3.867069 | 1.292969 | 0.344534 | 0.86398 |
| IF-upregulated | mmu04975 | Fat digestion and absorption | 40 | 2 | 1.237462 | 1.616211 | 0.352376 | 0.87026 |
| IF-upregulated | mmu05033 | Nicotine addiction | 40 | 2 | 1.237462 | 1.616211 | 0.352376 | 0.87026 |
| IF-upregulated | mmu00061 | Fatty acid biosynthesis | 14 | 1 | 0.433112 | 2.308873 | 0.356159 | 0.87299 |
| IF-upregulated | mmu04216 | Ferroptosis | 41 | 2 | 1.268399 | 1.576791 | 0.363647 | 0.88469 |
| IF-upregulated | mmu03022 | Basal transcription factors | 44 | 2 | 1.361208 | 1.469283 | 0.396987 | 0.89079 |
| IF-upregulated | mmu04270 | Vascular smooth muscle contraction | 129 | 5 | 3.990816 | 1.252877 | 0.369258 | 0.89079 |
| IF-upregulated | mmu00604 | Glycosphingolipid biosynthesis | 15 | 1 | 0.464048 | 2.154948 | 0.376111 | 0.89079 |
| IF-upregulated | mmu04976 | Bile secretion | 71 | 3 | 2.196495 | 1.365812 | 0.377354 | 0.89079 |
| IF-upregulated | mmu01100 | Metabolic pathways | 1351 | 44 | 41.795287 | 1.052750 | 0.378561 | 0.89079 |
| IF-upregulated | mmu04660 | T cell receptor signaling pathway | 101 | 4 | 3.124592 | 1.280167 | 0.381210 | 0.89079 |
| IF-upregulated | mmu03420 | Nucleotide excision repair | 43 | 2 | 1.330272 | 1.503452 | 0.385958 | 0.89079 |
| IF-upregulated | mmu04726 | Serotonergic synapse | 132 | 5 | 4.083625 | 1.224402 | 0.387819 | 0.89079 |
| IF-upregulated | mmu04915 | Estrogen signaling pathway | 133 | 5 | 4.114562 | 1.215196 | 0.394002 | 0.89079 |
| IF-upregulated | mmu00562 | Inositol phosphate metabolism | 73 | 3 | 2.258369 | 1.328393 | 0.394235 | 0.89079 |
| IF-upregulated | mmu00603 | Glycosphingolipid biosynthesis | 16 | 1 | 0.494985 | 2.020264 | 0.395447 | 0.89079 |
| IF-upregulated | mmu04971 | Gastric acid secretion | 74 | 3 | 2.289305 | 1.310441 | 0.402633 | 0.89079 |
| IF-upregulated | mmu04630 | JAK-STAT signaling pathway | 165 | 6 | 5.104532 | 1.175426 | 0.402778 | 0.89079 |

| | | | | | | | | |
|---|---|---|---|---|---|---|---|---|
| IF-upregulated | mmu04080 | Neuroactive ligand-receptor interaction | 289 | 10 | 8.940665 | 1.118485 | 0.404410 | 0.89079 |
| IF-upregulated | mmu05212 | Pancreatic cancer | 75 | 3 | 2.320242 | 1.292969 | 0.410997 | 0.89102 |
| IF-upregulated | mmu05146 | Amoebiasis | 106 | 4 | 3.279275 | 1.219782 | 0.416400 | 0.89102 |
| IF-upregulated | mmu00380 | Tryptophan metabolism | 46 | 2 | 1.423082 | 1.405401 | 0.418765 | 0.89102 |
| IF-upregulated | mmu05133 | Pertussis | 76 | 3 | 2.351178 | 1.275956 | 0.419325 | 0.89102 |
| IF-upregulated | mmu05164 | Influenza A | 168 | 6 | 5.197341 | 1.154436 | 0.419454 | 0.89102 |
| IF-upregulated | mmu01524 | Platinum drug resistance | 77 | 3 | 2.382115 | 1.259385 | 0.427615 | 0.89102 |
| IF-upregulated | mmu00565 | Ether lipid metabolism | 47 | 2 | 1.454018 | 1.375499 | 0.429504 | 0.89102 |
| IF-upregulated | mmu02010 | ABC transporters | 47 | 2 | 1.454018 | 1.375499 | 0.429504 | 0.89102 |
| IF-upregulated | mmu04371 | Apelin signaling pathway | 139 | 5 | 4.300181 | 1.162742 | 0.430946 | 0.89102 |
| IF-upregulated | mmu00770 | Pantothenate and CoA biosynthesis | 18 | 1 | 0.556858 | 1.795790 | 0.432347 | 0.89102 |
| IF-upregulated | mmu01230 | Biosynthesis of amino acids | 78 | 3 | 2.413051 | 1.243239 | 0.435863 | 0.89102 |
| IF-upregulated | mmu04931 | Insulin resistance | 109 | 4 | 3.372085 | 1.186210 | 0.437309 | 0.89102 |
| IF-upregulated | mmu00600 | Sphingolipid metabolism | 48 | 2 | 1.484955 | 1.346842 | 0.440138 | 0.89121 |
| IF-upregulated | mmu00670 | One carbon pool by folate | 19 | 1 | 0.587795 | 1.701275 | 0.449946 | 0.89989 |
| IF-upregulated | mmu01210 | 2-Oxocarboxylic acid metabolism | 19 | 1 | 0.587795 | 1.701275 | 0.449946 | 0.89989 |
| IF-upregulated | mmu00270 | Cysteine and methionine metabolism | 50 | 2 | 1.546828 | 1.292969 | 0.461076 | 0.90549 |
| IF-upregulated | mmu00330 | Arginine and proline metabolism | 50 | 2 | 1.546828 | 1.292969 | 0.461076 | 0.90549 |
| IF-upregulated | mmu00591 | Linoleic acid metabolism | 50 | 2 | 1.546828 | 1.292969 | 0.461076 | 0.90549 |
| IF-upregulated | mmu04724 | Glutamatergic synapse | 114 | 4 | 3.526767 | 1.134183 | 0.471660 | 0.92072 |
| IF-upregulated | mmu04350 | TGF-beta signaling pathway | 84 | 3 | 2.598671 | 1.154436 | 0.484337 | 0.92879 |
| IF-upregulated | mmu05014 | Amyotrophic lateral sclerosis (ALS) | 52 | 2 | 1.608701 | 1.243239 | 0.481551 | 0.92879 |
| IF-upregulated | mmu04146 | Peroxisome | 84 | 3 | 2.598671 | 1.154436 | 0.484337 | 0.92879 |
| IF-upregulated | mmu03320 | PPAR signaling pathway | 85 | 3 | 2.629607 | 1.140855 | 0.492227 | 0.93840 |
| IF-upregulated | mmu00514 | Other types of O-glycan biosynthesis | 22 | 1 | 0.680604 | 1.469283 | 0.499550 | 0.94135 |
| IF-upregulated | mmu04650 | Natural killer cell mediated cytotoxicity | 118 | 4 | 3.650514 | 1.095736 | 0.498576 | 0.94135 |
| IF-upregulated | mmu04020 | Calcium signaling pathway | 184 | 6 | 5.692326 | 1.054051 | 0.506824 | 0.94957 |
| IF-upregulated | mmu05167 | Kaposi sarcoma-associated herpesvirus infection | 217 | 7 | 6.713233 | 1.042717 | 0.509903 | 0.94988 |
| IF-upregulated | mmu04977 | Vitamin digestion and absorption | 24 | 1 | 0.742477 | 1.346842 | 0.530117 | 0.98192 |
| IF-upregulated | mmu00563 | Glycosylphosphatidylinositol (GPI)-anchor biosynthesis | 25 | 1 | 0.773414 | 1.292969 | 0.544696 | 0.98832 |
| IF-upregulated | mmu05163 | Human cytomegalovirus infection | 256 | 8 | 7.919758 | 1.010132 | 0.539827 | 0.98832 |
| IF-upregulated | mmu05134 | Legionellosis | 58 | 2 | 1.794320 | 1.114628 | 0.540010 | 0.98832 |
| IF-upregulated | mmu05032 | Morphine addiction | 92 | 3 | 2.846163 | 1.054051 | 0.545696 | 0.98832 |
| IF-downregulated | mmu00120 | Primary bile acid biosynthesis | 16 | 2 | 0.102477 | 19.516509 | 0.004560 | 1 |
| IF-downregulated | mmu01040 | Biosynthesis of unsaturated fatty acids | 32 | 2 | 0.204955 | 9.758255 | 0.017659 | 1 |
| IF-downregulated | mmu00430 | Taurine and hypotaurine metabolism | 11 | 1 | 0.070453 | 14.193825 | 0.068280 | 1 |
| IF-downregulated | mmu00230 | Purine metabolism | 179 | 3 | 1.146465 | 2.616739 | 0.106560 | 1 |
| IF-downregulated | mmu00052 | Galactose metabolism | 32 | 1 | 0.204955 | 4.879127 | 0.186168 | 1 |
| IF-downregulated | mmu04975 | Fat digestion and absorption | 40 | 1 | 0.256193 | 3.903302 | 0.227116 | 1 |
| IF-downregulated | mmu04072 | Phospholipase D signaling pathway | 147 | 2 | 0.941511 | 2.124246 | 0.242441 | 1 |
| IF-downregulated | mmu00240 | Pyrimidine metabolism | 101 | 1 | 0.646888 | 1.545862 | 0.479484 | 1 |
| IF-downregulated | mmu04630 | JAK-STAT signaling pathway | 165 | 4 | 1.056798 | 3.785020 | 0.020928 | 1 |
| IF-downregulated | mmu04064 | NF-kappa B signaling pathway | 104 | 3 | 0.666103 | 4.503810 | 0.028703 | 1 |
| IF-downregulated | mmu00380 | Tryptophan metabolism | 46 | 2 | 0.294622 | 6.788351 | 0.034816 | 1 |
| IF-downregulated | mmu04060 | Cytokine-cytokine receptor interaction | 296 | 5 | 1.895831 | 2.637366 | 0.040125 | 1 |
| IF-downregulated | mmu04210 | Apoptosis | 136 | 3 | 0.871057 | 3.444090 | 0.056106 | 1 |
| IF-downregulated | mmu04115 | p53 signaling pathway | 71 | 2 | 0.454743 | 4.398087 | 0.075607 | 1 |
| IF-downregulated | mmu04146 | Peroxisome | 84 | 2 | 0.538006 | 3.717430 | 0.100715 | 1 |

| | | | | | | | | |
|---|---|---|---|---|---|---|---|---|
| IF-downregulated | mmu03320 | PPAR signaling pathway | 85 | 2 | 0.544411 | 3.673696 | 0.102734 | 1 |
| IF-downregulated | mmu04080 | Neuroactive ligand-receptor interaction | 289 | 4 | 1.850997 | 2.160998 | 0.112977 | 1 |
| IF-downregulated | mmu04620 | Toll-like receptor signaling pathway | 99 | 2 | 0.634079 | 3.154183 | 0.132085 | 1 |
| IF-downregulated | mmu03060 | Protein export | 28 | 1 | 0.179335 | 5.576146 | 0.164903 | 1 |
| IF-downregulated | mmu04215 | Apoptosis | 32 | 1 | 0.204955 | 4.879127 | 0.186168 | 1 |
| IF-downregulated | mmu01100 | Metabolic pathways | 1351 | 11 | 8.652931 | 1.271246 | 0.238704 | 1 |
| IF-downregulated | mmu04010 | MAPK signaling pathway | 294 | 3 | 1.883021 | 1.593184 | 0.290955 | 1 |
| IF-downregulated | mmu00561 | Glycerolipid metabolism | 61 | 1 | 0.390695 | 2.559542 | 0.325230 | 1 |
| IF-downregulated | mmu04720 | Long-term potentiation | 67 | 1 | 0.429124 | 2.330329 | 0.350944 | 1 |
| IF-downregulated | mmu05016 | Huntington disease | 194 | 2 | 1.242538 | 1.609609 | 0.353969 | 1 |
| IF-downregulated | mmu04151 | PI3K-Akt signaling pathway | 358 | 3 | 2.292931 | 1.308369 | 0.404003 | 1 |
| IF-downregulated | mmu04012 | ErbB signaling pathway | 84 | 1 | 0.538006 | 1.858715 | 0.418690 | 1 |
| IF-downregulated | mmu04066 | HIF-1 signaling pathway | 105 | 1 | 0.672508 | 1.486972 | 0.492856 | 1 |
| IF-downregulated | mmu04668 | TNF signaling pathway | 110 | 1 | 0.704532 | 1.419383 | 0.509097 | 1 |
| IF-downregulated | mmu04722 | Neurotrophin signaling pathway | 121 | 1 | 0.774985 | 1.290348 | 0.543053 | 1 |
| IF-downregulated | mmu04071 | Sphingolipid signaling pathway | 122 | 1 | 0.781390 | 1.279771 | 0.546023 | 1 |
| IF-downregulated | mmu04110 | Cell cycle | 123 | 1 | 0.787795 | 1.269366 | 0.548975 | 1 |
| IF-downregulated | mmu04068 | FoxO signaling pathway | 132 | 1 | 0.845438 | 1.182819 | 0.574702 | 1 |
| IF-downregulated | mmu04978 | Mineral absorption | 45 | 2 | 0.288218 | 6.939203 | 0.033437 | 1 |
| IF-downregulated | mmu05217 | Basal cell carcinoma | 63 | 2 | 0.403505 | 4.956574 | 0.061353 | 1 |
| IF-downregulated | mmu04623 | Cytosolic DNA-sensing pathway | 64 | 2 | 0.409909 | 4.879127 | 0.063079 | 1 |
| IF-downregulated | mmu04934 | Cushing syndrome | 158 | 3 | 1.011964 | 2.964533 | 0.080130 | 1 |
| IF-downregulated | mmu01524 | Platinum drug resistance | 77 | 2 | 0.493172 | 4.055379 | 0.086920 | 1 |
| IF-downregulated | mmu03018 | RNA degradation | 83 | 2 | 0.531601 | 3.762219 | 0.098708 | 1 |
| IF-downregulated | mmu05202 | Transcriptional misregulation in cancer | 183 | 3 | 1.172085 | 2.559542 | 0.111944 | 1 |
| IF-downregulated | mmu04062 | Chemokine signaling pathway | 199 | 3 | 1.274562 | 2.353750 | 0.134490 | 1 |
| IF-downregulated | mmu04972 | Pancreatic secretion | 103 | 2 | 0.659698 | 3.031691 | 0.140798 | 1 |
| IF-downregulated | mmu04977 | Vitamin digestion and absorption | 24 | 1 | 0.153716 | 6.505503 | 0.143093 | 1 |
| IF-downregulated | mmu04928 | Parathyroid hormone synthesis, secretion and action | 107 | 2 | 0.685317 | 2.918357 | 0.149634 | 1 |
| IF-downregulated | mmu00790 | Folate biosynthesis | 26 | 1 | 0.166526 | 6.005080 | 0.154067 | 1 |
| IF-downregulated | mmu04625 | C-type lectin receptor signaling pathway | 112 | 2 | 0.717341 | 2.788073 | 0.160834 | 1 |
| IF-downregulated | mmu03020 | RNA polymerase | 30 | 1 | 0.192145 | 5.204403 | 0.175603 | 1 |
| IF-downregulated | mmu00500 | Starch and sucrose metabolism | 33 | 1 | 0.211360 | 4.731275 | 0.191401 | 1 |
| IF-downregulated | mmu05340 | Primary immunodeficiency | 36 | 1 | 0.230574 | 4.337002 | 0.206901 | 1 |
| IF-downregulated | mmu04973 | Carbohydrate digestion and absorption | 43 | 1 | 0.275408 | 3.630978 | 0.241945 | 1 |
| IF-downregulated | mmu03022 | Basal transcription factors | 44 | 1 | 0.281813 | 3.548456 | 0.246826 | 1 |
| IF-downregulated | mmu05321 | Inflammatory bowel disease (IBD) | 59 | 1 | 0.377885 | 2.646306 | 0.316439 | 1 |
| IF-downregulated | mmu04664 | Fc epsilon RI signaling pathway | 68 | 1 | 0.435529 | 2.296060 | 0.355135 | 1 |
| IF-downregulated | mmu04976 | Bile secretion | 71 | 1 | 0.454743 | 2.199043 | 0.367549 | 1 |
| IF-downregulated | mmu04260 | Cardiac muscle contraction | 78 | 1 | 0.499577 | 2.001693 | 0.395611 | 1 |
| IF-downregulated | mmu04512 | ECM-receptor interaction | 83 | 1 | 0.531601 | 1.881109 | 0.414904 | 1 |
| IF-downregulated | mmu04658 | Th1 and Th2 cell differentiation | 87 | 1 | 0.557221 | 1.794622 | 0.429902 | 1 |
| IF-downregulated | mmu01522 | Endocrine resistance | 93 | 1 | 0.595650 | 1.678840 | 0.451695 | 1 |
| IF-downregulated | mmu03015 | mRNA surveillance pathway | 96 | 1 | 0.614864 | 1.626376 | 0.462282 | 1 |
| IF-downregulated | mmu04659 | Th17 cell differentiation | 102 | 1 | 0.653293 | 1.530707 | 0.482859 | 1 |
| IF-downregulated | mmu03008 | Ribosome biogenesis in eukaryotes | 116 | 1 | 0.742961 | 1.345966 | 0.527914 | 1 |
| IF-downregulated | mmu04650 | Natural killer cell mediated cytotoxicity | 118 | 1 | 0.755770 | 1.323153 | 0.534028 | 1 |
| IF-downregulated | mmu04640 | Hematopoietic cell lineage | 95 | 3 | 0.608459 | 4.930487 | 0.022707 | 1 |

| | | | | | | | | |
|---|---|---|---|---|---|---|---|---|
| IF-downregulated | mmu05211 | Renal cell carcinoma | 68 | 2 | 0.435529 | 4.592120 | 0.070144 | 1 |
| IF-downregulated | mmu05221 | Acute myeloid leukemia | 69 | 2 | 0.441934 | 4.525567 | 0.071950 | 1 |
| IF-downregulated | mmu03010 | Ribosome | 175 | 3 | 1.120846 | 2.676550 | 0.101283 | 1 |
| IF-downregulated | mmu05210 | Colorectal cancer | 88 | 2 | 0.563625 | 3.548456 | 0.108857 | 1 |
| IF-downregulated | mmu00900 | Terpenoid backbone biosynthesis | 23 | 1 | 0.147311 | 6.788351 | 0.137554 | 1 |
| IF-downregulated | mmu05310 | Asthma | 25 | 1 | 0.160121 | 6.245283 | 0.148597 | 1 |
| IF-downregulated | mmu04611 | Platelet activation | 124 | 2 | 0.794199 | 2.518259 | 0.188298 | 1 |
| IF-downregulated | mmu05165 | Human papillomavirus infection | 370 | 4 | 2.369789 | 1.687914 | 0.211463 | 1 |
| IF-downregulated | mmu05216 | Thyroid cancer | 37 | 1 | 0.236979 | 4.219786 | 0.212003 | 1 |
| IF-downregulated | mmu05219 | Bladder cancer | 41 | 1 | 0.262598 | 3.808099 | 0.232091 | 1 |
| IF-downregulated | mmu05224 | Breast cancer | 147 | 2 | 0.941511 | 2.124246 | 0.242441 | 1 |
| IF-downregulated | mmu05226 | Gastric cancer | 150 | 2 | 0.960725 | 2.081761 | 0.249583 | 1 |
| IF-downregulated | mmu05206 | MicroRNAs in cancer | 281 | 3 | 1.799758 | 1.666890 | 0.268144 | 1 |
| IF-downregulated | mmu03460 | Fanconi anemia pathway | 51 | 1 | 0.326647 | 3.061413 | 0.280137 | 1 |
| IF-downregulated | mmu05225 | Hepatocellular carcinoma | 171 | 2 | 1.095227 | 1.826106 | 0.299664 | 1 |
| IF-downregulated | mmu05213 | Endometrial cancer | 58 | 1 | 0.371480 | 2.691932 | 0.312002 | 1 |
| IF-downregulated | mmu05152 | Tuberculosis | 178 | 2 | 1.140060 | 1.754293 | 0.316294 | 1 |
| IF-downregulated | mmu05223 | Non-small cell lung cancer | 66 | 1 | 0.422719 | 2.365638 | 0.346726 | 1 |
| IF-downregulated | mmu04510 | Focal adhesion | 199 | 2 | 1.274562 | 1.569167 | 0.365612 | 1 |
| IF-downregulated | mmu05214 | Glioma | 71 | 1 | 0.454743 | 2.199043 | 0.367549 | 1 |
| IF-downregulated | mmu05218 | Melanoma | 72 | 1 | 0.461148 | 2.168501 | 0.371635 | 1 |
| IF-downregulated | mmu05205 | Proteoglycans in cancer | 204 | 2 | 1.306586 | 1.530707 | 0.377180 | 1 |
| IF-downregulated | mmu05212 | Pancreatic cancer | 75 | 1 | 0.480363 | 2.081761 | 0.383737 | 1 |
| IF-downregulated | mmu05220 | Chronic myeloid leukemia | 76 | 1 | 0.486767 | 2.054369 | 0.387721 | 1 |
| IF-downregulated | mmu05132 | Salmonella infection | 78 | 1 | 0.499577 | 2.001693 | 0.395611 | 1 |
| IF-downregulated | mmu04610 | Complement and coagulation cascades | 88 | 1 | 0.563625 | 1.774228 | 0.433592 | 1 |
| IF-downregulated | mmu04742 | Taste transduction | 88 | 1 | 0.563625 | 1.774228 | 0.433592 | 1 |
| IF-downregulated | mmu05203 | Viral carcinogenesis | 231 | 2 | 1.479517 | 1.351793 | 0.438077 | 1 |
| IF-downregulated | mmu05200 | Pathways in cancer | 530 | 4 | 3.394562 | 1.178355 | 0.443524 | 1 |
| IF-downregulated | mmu05222 | Small cell lung cancer | 92 | 1 | 0.589245 | 1.697088 | 0.448120 | 1 |
| IF-downregulated | mmu05215 | Prostate cancer | 97 | 1 | 0.621269 | 1.609609 | 0.465767 | 1 |
| IF-downregulated | mmu04916 | Melanogenesis | 100 | 1 | 0.640483 | 1.561321 | 0.476087 | 1 |
| IF-downregulated | mmu05142 | Chagas disease (American trypanosomiasis) | 101 | 1 | 0.646888 | 1.545862 | 0.479484 | 1 |
| IF-downregulated | mmu05163 | Human cytomegalovirus infection | 256 | 2 | 1.639637 | 1.219782 | 0.491608 | 1 |
| IF-downregulated | mmu04670 | Leukocyte transendothelial migration | 115 | 1 | 0.736556 | 1.357670 | 0.524828 | 1 |
| IF-downregulated | mmu05166 | Human T-cell leukemia virus 1 infection | 282 | 2 | 1.806163 | 1.107320 | 0.543893 | 1 |
| IF-downregulated | mmu03040 | Spliceosome | 133 | 1 | 0.851843 | 1.173925 | 0.577470 | 1 |

**Supplemental Table 2.** 175 Gene enrichement

| source | term_name | term_id | adjusted_p_value |
|---|---|---|---|
| GO:BP | organonitrogen compound metabolic process | GO:1901564 | 1.77E-17 |
| GO:BP | cellular metabolic process | GO:0044237 | 2.00E-16 |
| GO:BP | metabolic process | GO:0008152 | 2.45E-16 |
| GO:BP | phosphate-containing compound metabolic process | GO:0006796 | 2.49E-15 |
| GO:CC | cytoplasm | GO:0005737 | 3.34E-15 |
| GO:BP | phosphorus metabolic process | GO:0006793 | 3.75E-15 |
| GO:BP | organic substance metabolic process | GO:0071704 | 9.29E-15 |
| GO:BP | cell surface receptor signaling pathway | GO:0007166 | 3.79E-13 |
| GO:BP | positive regulation of biological process | GO:0048518 | 4.35E-13 |
| GO:BP | phosphorylation | GO:0016310 | 6.75E-12 |
| GO:BP | positive regulation of cellular process | GO:0048522 | 1.75E-11 |
| GO:BP | protein metabolic process | GO:0019538 | 2.69E-11 |
| GO:BP | primary metabolic process | GO:0044238 | 1.12E-10 |
| GO:BP | regulation of phosphorylation | GO:0042325 | 3.24E-10 |
| GO:BP | regulation of molecular function | GO:0065009 | 3.35E-10 |
| GO:BP | positive regulation of cellular protein metabolic process | GO:0032270 | 3.54E-10 |
| GO:BP | cellular protein metabolic process | GO:0044267 | 4.28E-10 |
| GO:BP | nitrogen compound metabolic process | GO:0006807 | 6.53E-10 |
| GO:BP | positive regulation of protein metabolic process | GO:0051247 | 2.37E-09 |
| GO:BP | response to organic substance | GO:0010033 | 3.07E-09 |
| GO:BP | regulation of protein metabolic process | GO:0051246 | 3.99E-09 |
| GO:BP | regulation of phosphate metabolic process | GO:0019220 | 4.51E-09 |
| GO:BP | regulation of phosphorus metabolic process | GO:0051174 | 4.60E-09 |
| GO:BP | cellular process | GO:0009987 | 4.79E-09 |
| GO:BP | regulation of catalytic activity | GO:0050790 | 6.00E-09 |
| GO:BP | positive regulation of phosphate metabolic process | GO:0045937 | 6.16E-09 |
| GO:BP | positive regulation of phosphorus metabolic process | GO:0010562 | 6.16E-09 |
| GO:BP | intracellular signal transduction | GO:0035556 | 6.80E-09 |
| GO:BP | cell communication | GO:0007154 | 8.57E-09 |
| GO:BP | cellular protein modification process | GO:0006464 | 9.89E-09 |
| GO:BP | protein modification process | GO:0036211 | 9.89E-09 |
| GO:BP | positive regulation of metabolic process | GO:0009893 | 1.15E-08 |
| GO:BP | regulation of cellular protein metabolic process | GO:0032268 | 1.40E-08 |
| GO:BP | positive regulation of phosphorylation | GO:0042327 | 1.98E-08 |
| GO:BP | cellular response to stimulus | GO:0051716 | 2.13E-08 |
| GO:BP | regulation of cell communication | GO:0010646 | 3.06E-08 |
| GO:MF | protein binding | GO:0005515 | 3.57E-08 |
| GO:BP | regulation of signaling | GO:0023051 | 3.75E-08 |
| GO:BP | signaling | GO:0023052 | 4.02E-08 |
| GO:BP | positive regulation of protein modification process | GO:0031401 | 5.15E-08 |
| GO:BP | protein phosphorylation | GO:0006468 | 5.28E-08 |
| GO:BP | regulation of transferase activity | GO:0051338 | 5.72E-08 |
| GO:BP | regulation of signal transduction | GO:0009966 | 6.21E-08 |
| GO:BP | regulation of kinase activity | GO:0043549 | 6.86E-08 |
| GO:BP | positive regulation of cellular metabolic process | GO:0031325 | 8.66E-08 |
| GO:BP | signal transduction | GO:0007165 | 9.06E-08 |
| GO:BP | macromolecule modification | GO:0043412 | 9.64E-08 |
| GO:BP | regulation of protein phosphorylation | GO:0001932 | 9.78E-08 |
| GO:BP | response to stimulus | GO:0050896 | 1.09E-07 |
| GO:BP | catabolic process | GO:0009056 | 1.40E-07 |
| GO:BP | regulation of response to stimulus | GO:0048583 | 2.82E-07 |
| GO:BP | regulation of protein modification process | GO:0031399 | 3.14E-07 |
| GO:MF | enzyme binding | GO:0019899 | 3.32E-07 |
| GO:BP | positive regulation of protein phosphorylation | GO:0001934 | 3.41E-07 |
| GO:BP | positive regulation of nitrogen compound metabolic process | GO:0051173 | 4.83E-07 |
| GO:BP | positive regulation of macromolecule metabolic process | GO:0010604 | 5.71E-07 |

| | | | |
|---|---|---|---|
| GO:CC | intracellular | GO:0005622 | 6.39E-07 |
| GO:BP | cellular catabolic process | GO:0044248 | 1.13E-06 |
| GO:BP | response to nitrogen compound | GO:1901698 | 1.36E-06 |
| GO:BP | response to organonitrogen compound | GO:0010243 | 2.30E-06 |
| GO:MF | binding | GO:0005488 | 2.36E-06 |
| GO:BP | anatomical structure development | GO:0048856 | 4.00E-06 |
| GO:BP | organic substance catabolic process | GO:1901575 | 4.07E-06 |
| GO:MF | kinase binding | GO:0019900 | 4.36E-06 |
| GO:BP | positive regulation of cell communication | GO:0010647 | 4.91E-06 |
| GO:CC | membrane-bounded organelle | GO:0043227 | 5.05E-06 |
| GO:BP | regulation of intracellular signal transduction | GO:1902531 | 5.06E-06 |
| GO:BP | positive regulation of signaling | GO:0023056 | 5.42E-06 |
| GO:BP | regulation of multicellular organismal process | GO:0051239 | 5.47E-06 |
| GO:MF | catalytic activity | GO:0003824 | 5.64E-06 |
| GO:CC | intracellular membrane-bounded organelle | GO:0043231 | 6.46E-06 |
| GO:BP | cell-cell signaling | GO:0007267 | 7.58E-06 |
| GO:MF | identical protein binding | GO:0042802 | 7.95E-06 |
| GO:BP | small molecule metabolic process | GO:0044281 | 7.98E-06 |
| KEGG | Breast cancer | KEGG:05224 | 8.94E-06 |
| GO:BP | organonitrogen compound catabolic process | GO:1901565 | 9.06E-06 |
| GO:BP | positive regulation of signal transduction | GO:0009967 | 1.06E-05 |
| GO:CC | cytosol | GO:0005829 | 1.14E-05 |
| GO:MF | ubiquitin-like protein ligase binding | GO:0044389 | 1.15E-05 |
| GO:BP | organophosphate metabolic process | GO:0019637 | 1.27E-05 |
| GO:BP | cellular macromolecule metabolic process | GO:0044260 | 1.32E-05 |
| GO:BP | response to oxygen-containing compound | GO:1901700 | 1.32E-05 |
| GO:BP | response to external stimulus | GO:0009605 | 1.51E-05 |
| GO:BP | multicellular organism development | GO:0007275 | 1.57E-05 |
| GO:BP | positive regulation of intracellular signal transduction | GO:1902533 | 1.59E-05 |
| GO:BP | small molecule biosynthetic process | GO:0044283 | 1.71E-05 |
| GO:BP | system development | GO:0048731 | 2.41E-05 |
| GO:BP | macromolecule metabolic process | GO:0043170 | 3.06E-05 |
| GO:BP | developmental process | GO:0032502 | 3.31E-05 |
| GO:MF | ubiquitin protein ligase binding | GO:0031625 | 3.50E-05 |
| KEGG | Wnt signaling pathway | KEGG:04310 | 3.97E-05 |
| GO:BP | cellular response to organic substance | GO:0071310 | 4.96E-05 |
| GO:BP | cellular response to stress | GO:0033554 | 6.09E-05 |
| GO:BP | cell differentiation | GO:0030154 | 6.87E-05 |
| GO:BP | neuron projection development | GO:0031175 | 8.38E-05 |
| GO:BP | regulation of biological quality | GO:0065008 | 8.56E-05 |
| GO:MF | adenyl nucleotide binding | GO:0030554 | 8.56E-05 |
| GO:CC | organelle | GO:0043226 | 8.67E-05 |
| GO:BP | positive regulation of multicellular organismal process | GO:0051240 | 0.000100239 |
| GO:BP | regulation of protein serine/threonine kinase activity | GO:0071900 | 0.00010082 |
| GO:BP | lipid metabolic process | GO:0006629 | 0.000105069 |
| GO:BP | cellular lipid metabolic process | GO:0044255 | 0.000134197 |
| GO:BP | negative regulation of biological process | GO:0048519 | 0.000144788 |
| GO:BP | cellular developmental process | GO:0048869 | 0.000148374 |
| GO:BP | response to chemical | GO:0042221 | 0.00017147 |
| GO:BP | enzyme linked receptor protein signaling pathway | GO:0007167 | 0.000182742 |
| GO:BP | regulation of cell differentiation | GO:0045595 | 0.000198002 |
| GO:BP | regulation of metabolic process | GO:0019222 | 0.000201422 |
| GO:MF | protein kinase binding | GO:0019901 | 0.000201789 |
| GO:BP | neurogenesis | GO:0022008 | 0.000206288 |
| GO:BP | response to extracellular stimulus | GO:0009991 | 0.000216862 |
| GO:BP | neuron differentiation | GO:0030182 | 0.000218876 |
| GO:MF | small molecule binding | GO:0036094 | 0.000223653 |
| GO:BP | MAPK cascade | GO:0000165 | 0.000236429 |

| | | | |
|---|---|---|---|
| GO:BP | regulation of protein kinase activity | GO:0045859 | 0.000253455 |
| GO:BP | tube development | GO:0035295 | 0.000284406 |
| GO:BP | signal transduction by protein phosphorylation | GO:0023014 | 0.000298487 |
| GO:CC | intracellular organelle | GO:0043229 | 0.000300317 |
| GO:BP | neuron development | GO:0048666 | 0.000318101 |
| GO:BP | generation of neurons | GO:0048699 | 0.00032591 |
| GO:BP | animal organ development | GO:0048513 | 0.00032894 |
| GO:BP | anatomical structure morphogenesis | GO:0009653 | 0.000374698 |
| KEGG | Gastric cancer | KEGG:05226 | 0.000375732 |
| GO:BP | response to nutrient levels | GO:0031667 | 0.000377801 |
| KEGG | MAPK signaling pathway | KEGG:04010 | 0.000377966 |
| GO:BP | circulatory system development | GO:0072359 | 0.000497002 |
| GO:CC | cellular anatomical entity | GO:0110165 | 0.00053063 |
| GO:MF | purine nucleotide binding | GO:0017076 | 0.000583901 |
| GO:BP | cellular response to chemical stimulus | GO:0070887 | 0.000631298 |
| GO:BP | urogenital system development | GO:0001655 | 0.000644227 |
| GO:BP | cell death | GO:0008219 | 0.00067717 |
| GO:MF | adenyl ribonucleotide binding | GO:0032559 | 0.000678431 |
| GO:BP | positive regulation of stress-activated MAPK cascade | GO:0032874 | 0.000698563 |
| GO:BP | regulation of neuron projection development | GO:0010975 | 0.000739893 |
| GO:BP | response to endogenous stimulus | GO:0009719 | 0.000755612 |
| GO:BP | response to stress | GO:0006950 | 0.00075802 |
| GO:BP | negative regulation of cellular process | GO:0048523 | 0.000768887 |
| GO:BP | cell development | GO:0048468 | 0.00078123 |
| GO:BP | positive regulation of stress-activated protein kinase signaling cascade | GO:0070304 | 0.000788786 |
| GO:BP | regulation of multicellular organismal development | GO:2000026 | 0.000999674 |
| GO:BP | positive regulation of JNK cascade | GO:0046330 | 0.001058686 |
| GO:BP | regulation of MAPK cascade | GO:0043408 | 0.001059058 |
| GO:BP | regulation of cellular component organization | GO:0051128 | 0.001061538 |
| KEGG | Basal cell carcinoma | KEGG:05217 | 0.001073529 |
| GO:BP | positive regulation of catalytic activity | GO:0043085 | 0.001162002 |
| GO:BP | stress-activated MAPK cascade | GO:0051403 | 0.001194948 |
| GO:BP | positive regulation of molecular function | GO:0044093 | 0.00119767 |
| KEGG | MicroRNAs in cancer | KEGG:05206 | 0.001214476 |
| GO:BP | regulation of developmental process | GO:0050793 | 0.001381261 |
| GO:BP | nucleoside bisphosphate metabolic process | GO:0033865 | 0.001397546 |
| GO:BP | ribonucleoside bisphosphate metabolic process | GO:0033875 | 0.001397546 |
| GO:BP | purine nucleoside bisphosphate metabolic process | GO:0034032 | 0.001397546 |
| GO:BP | regulation of MAP kinase activity | GO:0043405 | 0.001447543 |
| GO:BP | regulation of neurogenesis | GO:0050767 | 0.001465771 |
| GO:BP | negative regulation of cell communication | GO:0010648 | 0.001574682 |
| GO:BP | negative regulation of signaling | GO:0023057 | 0.001651146 |
| GO:CC | protein-containing complex | GO:0032991 | 0.001724622 |
| GO:BP | nervous system development | GO:0007399 | 0.001726677 |
| KEGG | Hepatocellular carcinoma | KEGG:05225 | 0.001740536 |
| GO:BP | regulation of cell development | GO:0060284 | 0.001761788 |
| GO:MF | transferase activity | GO:0016740 | 0.001835445 |
| GO:BP | cell morphogenesis involved in differentiation | GO:0000904 | 0.001907126 |
| GO:BP | regulation of neuron differentiation | GO:0045664 | 0.00194971 |
| GO:BP | vasculature development | GO:0001944 | 0.002273178 |
| GO:BP | stress-activated protein kinase signaling cascade | GO:0031098 | 0.00228094 |
| GO:BP | positive regulation of MAPK cascade | GO:0043410 | 0.002324776 |
| GO:BP | cell part morphogenesis | GO:0032990 | 0.002415887 |
| KEGG | Colorectal cancer | KEGG:05210 | 0.002418114 |
| GO:BP | carbohydrate derivative metabolic process | GO:1901135 | 0.00256379 |
| GO:CC | whole membrane | GO:0098805 | 0.002656845 |
| GO:BP | tube morphogenesis | GO:0035239 | 0.002676394 |
| GO:BP | negative regulation of transferase activity | GO:0051348 | 0.002690687 |

| | | | |
|---|---|---|---|
| GO:CC | organelle membrane | GO:0031090 | 0.002722787 |
| KEGG | mTOR signaling pathway | KEGG:04150 | 0.002772567 |
| GO:BP | response to peptide | GO:1901652 | 0.002809954 |
| GO:BP | biological regulation | GO:0065007 | 0.002865086 |
| GO:BP | neuron projection morphogenesis | GO:0048812 | 0.002940686 |
| GO:BP | regulation of stress-activated MAPK cascade | GO:0032872 | 0.003161766 |
| GO:BP | apoptotic process | GO:0006915 | 0.003268234 |
| GO:BP | cell morphogenesis | GO:0000902 | 0.003269552 |
| GO:CC | mitochondrion | GO:0005739 | 0.003332497 |
| GO:BP | cell morphogenesis involved in neuron differentiation | GO:0048667 | 0.003503741 |
| GO:BP | regulation of stress-activated protein kinase signaling cascade | GO:0070302 | 0.003616101 |
| GO:BP | regulation of nervous system development | GO:0051960 | 0.003636751 |
| GO:MF | purine ribonucleotide binding | GO:0032555 | 0.003651423 |
| GO:BP | positive regulation of kinase activity | GO:0033674 | 0.00386046 |
| GO:BP | regulation of biological process | GO:0050789 | 0.003932049 |
| GO:BP | plasma membrane bounded cell projection morphogenesis | GO:0120039 | 0.004270427 |
| GO:BP | axon development | GO:0061564 | 0.004444326 |
| GO:MF | ribonucleotide binding | GO:0032553 | 0.004479922 |
| GO:BP | blood vessel development | GO:0001568 | 0.004600332 |
| GO:BP | thioester metabolic process | GO:0035383 | 0.004663289 |
| GO:BP | acyl-CoA metabolic process | GO:0006637 | 0.004663289 |
| GO:BP | cell projection morphogenesis | GO:0048858 | 0.004786881 |
| KEGG | Endometrial cancer | KEGG:05213 | 0.004820041 |
| GO:BP | programmed cell death | GO:0012501 | 0.00562473 |
| GO:BP | axonogenesis | GO:0007409 | 0.005692928 |
| GO:BP | regulation of cellular metabolic process | GO:0031323 | 0.005757859 |
| GO:MF | enzyme regulator activity | GO:0030234 | 0.005887322 |
| KEGG | Signaling pathways regulating pluripotency of stem cells | KEGG:04550 | 0.006321454 |
| GO:BP | protein localization | GO:0008104 | 0.006589643 |
| GO:MF | signaling receptor binding | GO:0005102 | 0.006932918 |
| GO:BP | regulation of anatomical structure morphogenesis | GO:0022603 | 0.007359021 |
| GO:BP | negative regulation of response to stimulus | GO:0048585 | 0.007429365 |
| GO:MF | carbohydrate derivative binding | GO:0097367 | 0.007853648 |
| KEGG | Sphingolipid signaling pathway | KEGG:04071 | 0.00786561 |
| GO:BP | regulation of cellular process | GO:0050794 | 0.00810422 |
| GO:BP | organophosphate biosynthetic process | GO:0090407 | 0.00846998 |
| GO:BP | proteasomal protein catabolic process | GO:0010498 | 0.008708672 |
| GO:MF | kinase activity | GO:0016301 | 0.009137259 |
| GO:BP | positive regulation of response to stimulus | GO:0048584 | 0.009145212 |
| GO:MF | protein phosphatase binding | GO:0019903 | 0.009356589 |
| GO:BP | cell surface receptor signaling pathway involved in cell-cell signaling | GO:1905114 | 0.009356992 |
| GO:MF | nucleotide binding | GO:0000166 | 0.009460669 |
| GO:MF | nucleoside phosphate binding | GO:1901265 | 0.009460669 |
| GO:MF | transferase activity, transferring phosphorus-containing groups | GO:0016772 | 0.009563118 |
| GO:BP | positive regulation of cell population proliferation | GO:0008284 | 0.010506296 |
| GO:CC | perinuclear region of cytoplasm | GO:0048471 | 0.012142934 |
| GO:BP | regulation of response to stress | GO:0080134 | 0.012293944 |
| GO:BP | organic substance transport | GO:0071702 | 0.01237519 |
| GO:MF | molecular function regulator | GO:0098772 | 0.012539892 |
| GO:BP | protein transport | GO:0015031 | 0.012869969 |
| GO:BP | protein catabolic process | GO:0030163 | 0.013404346 |
| GO:BP | nitrogen compound transport | GO:0071705 | 0.013527641 |
| GO:MF | interleukin-11 receptor activity | GO:0004921 | 0.01378866 |
| GO:MF | interleukin-11 binding | GO:0019970 | 0.01378866 |
| GO:BP | positive regulation of protein kinase activity | GO:0045860 | 0.014558633 |
| GO:BP | negative regulation of developmental process | GO:0051093 | 0.014809186 |
| GO:BP | regulation of primary metabolic process | GO:0080090 | 0.014834312 |
| GO:BP | plasma membrane bounded cell projection organization | GO:0120036 | 0.015331687 |

| | | | |
|---|---|---|---|
| GO:BP | establishment of protein localization | GO:0045184 | 0.016003536 |
| GO:BP | cellular component morphogenesis | GO:0032989 | 0.016036937 |
| GO:BP | chemotaxis | GO:0006935 | 0.016656361 |
| GO:BP | regulation of macromolecule metabolic process | GO:0060255 | 0.016852107 |
| KEGG | Hippo signaling pathway | KEGG:04390 | 0.017550889 |
| GO:BP | kidney development | GO:0001822 | 0.017663071 |
| GO:BP | taxis | GO:0042330 | 0.017829288 |
| GO:BP | negative regulation of cell development | GO:0010721 | 0.017953566 |
| KEGG | Pathways in cancer | KEGG:05200 | 0.018592194 |
| GO:BP | purine nucleotide metabolic process | GO:0006163 | 0.019070131 |
| GO:BP | regulation of JNK cascade | GO:0046328 | 0.019089128 |
| GO:BP | anatomical structure formation involved in morphogenesis | GO:0048646 | 0.019620602 |
| GO:BP | regulation of cell death | GO:0010941 | 0.020575356 |
| GO:BP | response to nutrient | GO:0007584 | 0.020972901 |
| GO:MF | phosphotransferase activity, alcohol group as acceptor | GO:0016773 | 0.022506843 |
| GO:BP | positive regulation of developmental process | GO:0051094 | 0.023099062 |
| GO:BP | macromolecule localization | GO:0033036 | 0.023284473 |
| GO:BP | regulation of cell morphogenesis involved in differentiation | GO:0010769 | 0.02371427 |
| KEGG | Thyroid cancer | KEGG:05216 | 0.024119095 |
| KEGG | Melanoma | KEGG:05218 | 0.024312405 |
| GO:BP | positive regulation of cellular component organization | GO:0051130 | 0.024423462 |
| GO:BP | multicellular organismal process | GO:0032501 | 0.024834512 |
| GO:BP | peptide transport | GO:0015833 | 0.025248956 |
| GO:BP | renal system development | GO:0072001 | 0.025285013 |
| GO:BP | cellular response to starvation | GO:0009267 | 0.025826065 |
| GO:BP | positive regulation of transferase activity | GO:0051347 | 0.026574086 |
| GO:BP | cell projection organization | GO:0030030 | 0.026632692 |
| KEGG | Glioma | KEGG:05214 | 0.026879281 |
| GO:BP | locomotion | GO:0040011 | 0.028102999 |
| GO:MF | protein domain specific binding | GO:0019904 | 0.028143125 |
| GO:BP | phospholipid metabolic process | GO:0006644 | 0.028776908 |
| GO:BP | negative regulation of multicellular organismal process | GO:0051241 | 0.029824215 |
| GO:BP | response to starvation | GO:0042594 | 0.030213804 |
| GO:BP | muscle cell proliferation | GO:0033002 | 0.032015057 |
| GO:BP | response to peptide hormone | GO:0043434 | 0.032565885 |
| GO:BP | JNK cascade | GO:0007254 | 0.03301092 |
| GO:BP | negative regulation of kinase activity | GO:0033673 | 0.033273819 |
| GO:BP | blood vessel morphogenesis | GO:0048514 | 0.034382145 |
| GO:MF | anion binding | GO:0043168 | 0.03454616 |
| GO:BP | regulation of cellular localization | GO:0060341 | 0.035145756 |
| GO:BP | purine ribonucleotide metabolic process | GO:0009150 | 0.035682861 |
| GO:BP | N-glycan processing | GO:0006491 | 0.0357441 |
| GO:BP | regulation of localization | GO:0032879 | 0.037171458 |
| GO:BP | behavior | GO:0007610 | 0.037503217 |
| GO:MF | kinase regulator activity | GO:0019207 | 0.037868138 |
| GO:BP | amide transport | GO:0042886 | 0.038106681 |
| GO:BP | signal release | GO:0023061 | 0.038204247 |
| GO:BP | regulation of plasma membrane bounded cell projection organization | GO:0120035 | 0.038564501 |
| GO:BP | establishment of localization in cell | GO:0051649 | 0.039231995 |
| GO:BP | Wnt signaling pathway | GO:0016055 | 0.040358016 |
| KEGG | Melanogenesis | KEGG:04916 | 0.040892782 |
| GO:BP | regulation of nitrogen compound metabolic process | GO:0051171 | 0.041005226 |
| GO:BP | cell-cell signaling by wnt | GO:0198738 | 0.04264555 |
| GO:CC | bounding membrane of organelle | GO:0098588 | 0.043173715 |
| GO:BP | transmembrane receptor protein tyrosine kinase signaling pathway | GO:0007169 | 0.044790669 |
| GO:BP | regulation of cell projection organization | GO:0031344 | 0.046453547 |
| GO:BP | regulation of cell cycle process | GO:0010564 | 0.046794806 |
| GO:MF | amidinotransferase activity | GO:0015067 | 0.049013806 |

| | | | |
|---|---|---|---|
| GO:MF | glycine amidinotransferase activity | GO:0015068 | 0.049013806 |
| GO:CC | proteasome accessory complex | GO:0022624 | 0.04968191 |
| GO:BP | tissue development | GO:0009888 | 0.04968771 |

### **Supplementary Table 3.** PICRUSt - bacteria pathway

| Taxon | AD+Ad Libitum | AD+IF | Fold change |
|---|---|---|---|
| Apoptosis | 3.72E-05 | 0.000129531 | 3.4800679 |
| RIG-I-like receptor signaling pathway | 3.04E-05 | 0.000102288 | 3.368136111 |
| Staphylococcus aureus infection | 0.000268386 | 0.000736115 | 2.742750206 |
| Transcription related proteins | 6.48E-05 | 0.000173285 | 2.674175805 |
| Ethylbenzene degradation | 0.00049793 | 0.001036725 | 2.082071306 |
| Carbohydrate digestion and absorption | 7.01E-05 | 0.000130263 | 1.859096909 |
| Ion channels | 0.00030252 | 0.00048315 | 1.597084188 |
| Limonene and pinene degradation | 0.001017631 | 0.001605489 | 1.577673751 |
| Bisphenol degradation | 0.001080538 | 0.001666033 | 1.541854128 |
| Naphthalene degradation | 0.001395978 | 0.002130401 | 1.526098613 |
| Chlorocyclohexane and chlorobenzene degradation | 0.000114155 | 0.000171982 | 1.506566215 |
| Linoleic acid metabolism | 0.000930052 | 0.001375035 | 1.478449388 |
| Aminobenzoate degradation | 0.001475532 | 0.002173106 | 1.472760853 |
| Xylene degradation | 0.000578081 | 0.000797648 | 1.379820426 |
| Tetracycline biosynthesis | 0.000916395 | 0.001212125 | 1.322709474 |
| Dioxin degradation | 0.000639268 | 0.000842639 | 1.318130919 |
| Secondary bile acid biosynthesis | 0.000349536 | 0.00045576 | 1.303901231 |
| Primary bile acid biosynthesis | 0.000350236 | 0.000456138 | 1.302373166 |
| Benzoate degradation | 0.002544155 | 0.003274982 | 1.287257482 |
| Fatty acid biosynthesis | 0.003661199 | 0.004594212 | 1.254838069 |
| Toluene degradation | 0.000587846 | 0.000733373 | 1.247558873 |
| Stilbenoid, diarylheptanoid and gingerol biosynthesis | 0.000166079 | 0.000205022 | 1.23448741 |
| Phosphonate and phosphinate metabolism | 0.000404178 | 0.000496193 | 1.227658141 |
| Tyrosine metabolism | 0.003674663 | 0.004490704 | 1.222072425 |
| Chloroalkane and chloroalkene degradation | 0.002329967 | 0.002833886 | 1.216277606 |
| Fructose and mannose metabolism | 0.01312904 | 0.015893483 | 1.210559359 |
| Caprolactam degradation | 0.000221999 | 0.00026783 | 1.206445914 |
| Glycosyltransferases | 0.003076886 | 0.003647379 | 1.185412604 |
| D-Arginine and D-ornithine metabolism | 0.000207631 | 0.000245823 | 1.183940793 |
| Ubiquitin system | 3.00E-05 | 3.53E-05 | 1.177452718 |
| Others | 0.011150585 | 0.012968621 | 1.163043945 |
| Proximal tubule bicarbonate reclamation | 8.21E-05 | 9.52E-05 | 1.160113414 |
| D-Alanine metabolism | 0.001359329 | 0.001576493 | 1.159757631 |
| Biosynthesis of unsaturated fatty acids | 0.000960253 | 0.001112513 | 1.158563312 |
| Lipoic acid metabolism | 0.000291325 | 0.000333665 | 1.145335651 |
| Glutathione metabolism | 0.00185165 | 0.002103811 | 1.13618189 |
| Butanoate metabolism | 0.006146656 | 0.006938721 | 1.128861262 |
| Renal cell carcinoma | 0.000227788 | 0.000255791 | 1.122935568 |
| Replication, recombination and repair proteins | 0.00807101 | 0.009041258 | 1.120213887 |
| Glycerolipid metabolism | 0.004206863 | 0.004710495 | 1.119716812 |
| Phosphotransferase system (PTS) | 0.010738944 | 0.011921122 | 1.110083285 |
| Lipid biosynthesis proteins | 0.004805917 | 0.005315644 | 1.106062345 |
| Polycyclic aromatic hydrocarbon degradation | 0.001335356 | 0.001474835 | 1.104450954 |
| Synthesis and degradation of ketone bodies | 0.000615214 | 0.000678795 | 1.103348889 |
| Metabolism of cofactors and vitamins | 0.000850261 | 0.00093518 | 1.09987416 |
| Starch and sucrose metabolism | 0.009024073 | 0.009922399 | 1.099547704 |
| Primary immunodeficiency | 0.00045517 | 0.000492489 | 1.081988822 |
| Drug metabolism - other enzymes | 0.002978985 | 0.003217726 | 1.080141608 |
| RNA polymerase | 0.001773554 | 0.001903881 | 1.073483742 |
| Phosphatidylinositol signaling system | 0.000999727 | 0.001072832 | 1.073125233 |
| Glycolysis / Gluconeogenesis | 0.011424415 | 0.012245134 | 1.071839036 |
| Signal transduction mechanisms | 0.005044548 | 0.00538986 | 1.068452617 |
| Pyruvate metabolism | 0.009612092 | 0.010228141 | 1.064091045 |
| Purine metabolism | 0.02242686 | 0.02379596 | 1.061047354 |
| Taurine and hypotaurine metabolism | 0.001307486 | 0.001384912 | 1.059217336 |
| Inositol phosphate metabolism | 0.001006972 | 0.001064156 | 1.056787933 |
| Butirosin and neomycin biosynthesis | 0.000683087 | 0.000720819 | 1.055238463 |
| RNA transport | 0.001651454 | 0.001739879 | 1.053544193 |
| Ubiquinone and other terpenoid-quinone biosynthesis | 0.001466624 | 0.001543363 | 1.052323594 |
| Other ion-coupled transporters | 0.011867135 | 0.012465204 | 1.050397064 |
| Sphingolipid metabolism | 0.00160853 | 0.001678558 | 1.043535378 |
| Glycosphingolipid biosynthesis - globo series | 0.000849984 | 0.000886276 | 1.042697737 |
| Sulfur metabolism | 0.00239099 | 0.002487709 | 1.040451325 |
| Amino sugar and nucleotide sugar metabolism | 0.017791494 | 0.018461299 | 1.037647485 |
| Transcription factors | 0.017097842 | 0.017736299 | 1.037341381 |
| Streptomycin biosynthesis | 0.003128863 | 0.003240562 | 1.03569965 |
| Lysine biosynthesis | 0.007292646 | 0.007545652 | 1.034693298 |
| Prenyltransferases | 0.003008891 | 0.00311273 | 1.034510462 |
| Function unknown | 0.012641361 | 0.013053618 | 1.032611709 |
| Lipid metabolism | 0.001248838 | 0.001287018 | 1.030572543 |
| Selenocompound metabolism | 0.003971885 | 0.004085555 | 1.028618774 |

| | | | |
|---|---|---|---|
| Restriction enzyme | 0.002182541 | 0.002244334 | 1.028312361 |
| Cytoskeleton proteins | 0.003815647 | 0.003923439 | 1.028250215 |
| Nucleotide excision repair | 0.003979519 | 0.004081939 | 1.025736669 |
| Chromosome | 0.015467193 | 0.015837846 | 1.023963784 |
| Amino acid metabolism | 0.002291305 | 0.002346037 | 1.023886956 |
| Glycerophospholipid metabolism | 0.00546231 | 0.005590683 | 1.023501724 |
| Propanoate metabolism | 0.005060159 | 0.005178695 | 1.023425325 |
| Cell cycle - Caulobacter | 0.005090957 | 0.00519272 | 1.019989052 |
| Pyrimidine metabolism | 0.018666113 | 0.019023813 | 1.019163066 |
| Protein kinases | 0.002708765 | 0.002745628 | 1.01360877 |
| D-Glutamine and D-glutamate metabolism | 0.001464072 | 0.001483975 | 1.013594055 |
| DNA repair and recombination proteins | 0.028396151 | 0.028771143 | 1.013205737 |
| Pentose phosphate pathway | 0.009327372 | 0.009450345 | 1.013184072 |
| Mismatch repair | 0.008282882 | 0.008388641 | 1.012768459 |
| Cysteine and methionine metabolism | 0.009441721 | 0.009534755 | 1.009853501 |
| Galactose metabolism | 0.007713002 | 0.007788093 | 1.009735716 |
| DNA replication proteins | 0.012447459 | 0.012521004 | 1.005908388 |
| DNA replication | 0.00701501 | 0.007054586 | 1.005641639 |
| Peptidoglycan biosynthesis | 0.008462245 | 0.008503297 | 1.004851251 |
| Ribosome | 0.024416447 | 0.024530177 | 1.004657891 |
| Ribosome Biogenesis | 0.014349917 | 0.014395698 | 1.003190337 |
| Translation factors | 0.005472773 | 0.005483241 | 1.001912693 |
| Ribosome biogenesis in eukaryotes | 0.000505432 | 0.000505164 | 0.999469979 |
| Aminoacyl-tRNA biosynthesis | 0.01293187 | 0.012924884 | 0.999459776 |
| Transporters | 0.075486868 | 0.075270035 | 0.997127529 |
| One carbon pool by folate | 0.005733464 | 0.005716692 | 0.997074677 |
| Inorganic ion transport and metabolism | 0.001763644 | 0.001756214 | 0.995787151 |
| Terpenoid backbone biosynthesis | 0.00604699 | 0.00602111 | 0.995720113 |
| Pentose and glucuronate interconversions | 0.004781918 | 0.004757675 | 0.994930407 |
| Carbon fixation in photosynthetic organisms | 0.006497496 | 0.006461588 | 0.994473587 |
| Homologous recombination | 0.009154172 | 0.009102425 | 0.994347166 |
| Type II diabetes mellitus | 0.000473686 | 0.000470894 | 0.994105591 |
| Chaperones and folding catalysts | 0.008888025 | 0.008830457 | 0.993523004 |
| Peptidases | 0.020245067 | 0.020111017 | 0.99337864 |
| Thiamine metabolism | 0.005022561 | 0.004977018 | 0.990932165 |
| Nucleotide metabolism | 0.000826628 | 0.000817712 | 0.989214124 |
| Chagas disease (American trypanosomiasis) | 1.70E-05 | 1.69E-05 | 0.988660078 |
| Methane metabolism | 0.012389868 | 0.012231264 | 0.987198938 |
| Translation proteins | 0.009052898 | 0.008934554 | 0.986927557 |
| Type I diabetes mellitus | 0.000497522 | 0.000490009 | 0.984899859 |
| Amino acid related enzymes | 0.015090073 | 0.014840191 | 0.983440592 |
| Zeatin biosynthesis | 0.000480438 | 0.000472478 | 0.983433092 |
| African trypanosomiasis | 1.93E-05 | 1.89E-05 | 0.981600865 |
| RNA degradation | 0.00412531 | 0.00404747 | 0.981130996 |
| Base excision repair | 0.00470511 | 0.004600983 | 0.977869291 |
| Pantothenate and CoA biosynthesis | 0.005248653 | 0.005114546 | 0.974449256 |
| Tuberculosis | 0.001561191 | 0.001520582 | 0.97398824 |
| Ascorbate and aldarate metabolism | 0.001020296 | 0.000992317 | 0.972577321 |
| Alanine, aspartate and glutamate metabolism | 0.010518845 | 0.010226518 | 0.97220915 |
| Polyketide sugar unit biosynthesis | 0.001989636 | 0.001933743 | 0.971908151 |
| Isoquinoline alkaloid biosynthesis | 0.000537209 | 0.00052131 | 0.970404232 |
| Flavone and flavonol biosynthesis | 8.90E-05 | 8.63E-05 | 0.969720354 |
| Glycine, serine and threonine metabolism | 0.007467303 | 0.007210195 | 0.965568928 |
| Metabolism of xenobiotics by cytochrome P450 | 0.000193694 | 0.000186973 | 0.965298452 |
| Other glycan degradation | 0.002241135 | 0.002163018 | 0.965144098 |
| Carbon fixation pathways in prokaryotes | 0.008456682 | 0.008157528 | 0.964625121 |
| Bacterial toxins | 0.001310604 | 0.001261661 | 0.962656276 |
| Protein export | 0.006058009 | 0.005826906 | 0.961851706 |
| Citrate cycle (TCA cycle) | 0.004681909 | 0.004490215 | 0.959056495 |
| Transcription machinery | 0.008191381 | 0.007844978 | 0.957711305 |
| Cyanoamino acid metabolism | 0.002265529 | 0.002166233 | 0.956170615 |
| General function prediction only | 0.037400235 | 0.035723966 | 0.955180251 |
| Biosynthesis of vancomycin group antibiotics | 0.000589811 | 0.000563287 | 0.955029625 |
| Retinol metabolism | 0.000204577 | 0.000195352 | 0.954907977 |
| Folate biosynthesis | 0.003515167 | 0.003355196 | 0.954491324 |
| Prion diseases | 6.45E-05 | 6.15E-05 | 0.953713481 |
| Carbohydrate metabolism | 0.001867552 | 0.001781092 | 0.953704162 |
| Vibrio cholerae pathogenic cycle | 0.000691911 | 0.000659334 | 0.952917505 |
| Secretion system | 0.012330721 | 0.011736518 | 0.951811136 |
| Valine, leucine and isoleucine biosynthesis | 0.005658214 | 0.005381429 | 0.95108278 |
| Sulfur relay system | 0.002620878 | 0.002488123 | 0.949347358 |
| Glycosaminoglycan degradation | 0.000345456 | 0.000327423 | 0.947798803 |
| ABC transporters | 0.033452927 | 0.031664003 | 0.946524124 |
| Photosynthesis | 0.004662016 | 0.004405047 | 0.944880329 |

| | | | |
|---|---|---|---|
| Photosynthesis proteins | 0.00474005 | 0.004473541 | 0.943775146 |
| beta-Lactam resistance | 9.53E-05 | 8.99E-05 | 0.943663501 |
| Valine, leucine and isoleucine degradation | 0.002161498 | 0.002037172 | 0.942481436 |
| Nicotinate and nicotinamide metabolism | 0.004501601 | 0.004232123 | 0.940137292 |
| Membrane and intracellular structural molecules | 0.003673461 | 0.003449953 | 0.939156056 |
| Glycosphingolipid biosynthesis - ganglio series | 0.00021401 | 0.00020094 | 0.938930574 |
| Bacterial secretion system | 0.005357113 | 0.005012728 | 0.935714453 |
| Nitrogen metabolism | 0.006719473 | 0.006286389 | 0.935547918 |
| C5-Branched dibasic acid metabolism | 0.002208467 | 0.002060113 | 0.93282455 |
| Pores ion channels | 0.001927562 | 0.001797569 | 0.932560848 |
| Lysosome | 0.000910152 | 0.000846625 | 0.930201359 |
| Arginine and proline metabolism | 0.010960737 | 0.010171378 | 0.927983062 |
| PPAR signaling pathway | 0.000860968 | 0.000798282 | 0.927192038 |
| Drug metabolism - cytochrome P450 | 0.000206765 | 0.00019133 | 0.925352123 |
| Oxidative phosphorylation | 0.010305201 | 0.009529671 | 0.924743783 |
| Fatty acid metabolism | 0.002281363 | 0.002108316 | 0.924147722 |
| Two-component system | 0.014847119 | 0.013655851 | 0.919764387 |
| Plant-pathogen interaction | 0.001636626 | 0.001500647 | 0.916914952 |
| Riboflavin metabolism | 0.00212491 | 0.001940126 | 0.913039128 |
| Porphyrin and chlorophyll metabolism | 0.007076052 | 0.006450069 | 0.91153502 |
| Protein folding and associated processing | 0.007240533 | 0.006597591 | 0.911202454 |
| Glutamatergic synapse | 0.001088 | 0.000989568 | 0.909529096 |
| Alzheimer's disease | 0.000597654 | 0.000543068 | 0.908666166 |
| Cell motility and secretion | 0.001393897 | 0.001266397 | 0.908529566 |
| Vitamin B6 metabolism | 0.001580249 | 0.001435417 | 0.908348083 |
| Pathways in cancer | 0.000631433 | 0.000568892 | 0.900954125 |
| Energy metabolism | 0.007713228 | 0.006922182 | 0.897442976 |
| Lipopolysaccharide biosynthesis proteins | 0.002360102 | 0.002113475 | 0.8955013 |
| Other transporters | 0.002620856 | 0.002343006 | 0.893985109 |
| Phenylalanine metabolism | 0.002114422 | 0.001887699 | 0.892773085 |
| Tropane, piperidine and pyridine alkaloid biosynthesis | 0.001041362 | 0.000922029 | 0.885406923 |
| Sporulation | 0.005705097 | 0.00503798 | 0.883066524 |
| Insulin signaling pathway | 0.000732273 | 0.000645851 | 0.881981248 |
| Peroxisome | 0.001370497 | 0.001205539 | 0.879636549 |
| Biotin metabolism | 0.001263383 | 0.001107667 | 0.876746589 |
| beta-Alanine metabolism | 0.00144129 | 0.001262868 | 0.876207123 |
| Amyotrophic lateral sclerosis (ALS) | 0.000175574 | 0.000152004 | 0.865754959 |
| Biosynthesis of siderophore group nonribosomal peptides | 0.000259549 | 0.00022446 | 0.8648082 |
| Novobiocin biosynthesis | 0.001421504 | 0.001227714 | 0.863672441 |
| Histidine metabolism | 0.006602349 | 0.005690062 | 0.861823806 |
| Atrazine degradation | 0.000205272 | 0.000176231 | 0.858522435 |
| MAPK signaling pathway - yeast | 0.000235665 | 0.000201835 | 0.856448799 |
| Biosynthesis of ansamycins | 0.001477183 | 0.001255644 | 0.850026092 |
| Phenylpropanoid biosynthesis | 0.00110067 | 0.000934285 | 0.848833277 |
| Protein digestion and absorption | 5.27E-05 | 4.45E-05 | 0.844045624 |
| Steroid hormone biosynthesis | 7.86E-05 | 6.59E-05 | 0.839408313 |
| Styrene degradation | 0.000210431 | 0.00017656 | 0.839040345 |
| Flagellar assembly | 0.005246599 | 0.004392825 | 0.837270904 |
| Glyoxylate and dicarboxylate metabolism | 0.005400238 | 0.004514805 | 0.836038108 |
| Tryptophan metabolism | 0.001654369 | 0.001378583 | 0.833298406 |
| Phenylalanine, tyrosine and tryptophan biosynthesis | 0.007590469 | 0.006295875 | 0.829444828 |
| Lysine degradation | 0.001332901 | 0.001098965 | 0.824490463 |
| Flavonoid biosynthesis | 0.00017771 | 0.000146198 | 0.822678604 |
| Cellular antigens | 0.00053747 | 0.000440827 | 0.820188176 |
| Bacterial chemotaxis | 0.005864567 | 0.004807315 | 0.81972204 |
| Various types of N-glycan biosynthesis | 1.32E-05 | 1.08E-05 | 0.817213964 |
| Bacterial motility proteins | 0.01183909 | 0.009605284 | 0.811319472 |
| Nitrotoluene degradation | 0.000859685 | 0.00069288 | 0.805970229 |
| Cell division | 0.00058002 | 0.000466342 | 0.804009005 |
| Electron transfer carriers | 0.000295369 | 0.000235815 | 0.798375104 |
| Pertussis | 3.64E-05 | 2.90E-05 | 0.795786052 |
| Penicillin and cephalosporin biosynthesis | 9.73E-05 | 7.70E-05 | 0.791701742 |
| Epithelial cell signaling in Helicobacter pylori infection | 0.000862997 | 0.00068211 | 0.790396866 |
| Proteasome | 0.000452012 | 0.000353555 | 0.782181524 |
| N-Glycan biosynthesis | 0.000220321 | 0.000171871 | 0.780094967 |
| Protein processing in endoplasmic reticulum | 0.000631981 | 0.000492954 | 0.780014638 |
| Adipocytokine signaling pathway | 0.000420949 | 0.000327891 | 0.7789318 |
| NOD-like receptor signaling pathway | 0.000405873 | 0.000315963 | 0.778477832 |
| Progesterone-mediated oocyte maturation | 0.000402827 | 0.000312719 | 0.776309936 |
| Antigen processing and presentation | 0.000402827 | 0.000312719 | 0.776309936 |
| Prostate cancer | 0.00040318 | 0.000312928 | 0.776149521 |
| Lipopolysaccharide biosynthesis | 0.001481292 | 0.001129187 | 0.762298914 |
| Amoebiasis | 0.000139837 | 0.000106372 | 0.760687675 |
| Biosynthesis and biodegradation of secondary metabolites | 0.000435805 | 0.000328937 | 0.754781464 |

| | | | |
|---|---|---|---|
| Glycan biosynthesis and metabolism | 0.000247653 | 0.000170882 | 0.690005979 |
| Arachidonic acid metabolism | 0.000251324 | 0.000171401 | 0.681992172 |
| Geraniol degradation | 0.000168057 | 0.000114201 | 0.679534919 |
| Meiosis - yeast | 7.96E-05 | 5.40E-05 | 0.678008173 |
| Huntington's disease | 0.000233741 | 0.000157291 | 0.672928001 |
| Bacterial invasion of epithelial cells | 1.94E-05 | 1.30E-05 | 0.66830569 |
| Mineral absorption | 4.26E-05 | 2.75E-05 | 0.645675636 |
| Germination | 0.000388403 | 0.000247254 | 0.636589785 |
| Basal transcription factors | 1.43E-05 | 8.22E-06 | 0.574619634 |
| Others | 3.54E-05 | 1.42E-05 | 0.402052995 |
| alpha-Linolenic acid metabolism | 2.59E-05 | 1.03E-05 | 0.399056914 |
| Non-homologous end-joining | 3.14E-05 | 1.23E-05 | 0.392166013 |
| Parkinson's disease | 3.68E-05 | 9.10E-06 | 0.247393319 |
| Cardiac muscle contraction | 3.50E-05 | 8.37E-06 | 0.238897513 |

**Supplementary Table 4.** Primer sequences used for RT-qPCR

| | Forward Primers (5'-3') | Reverse Primers (5'-3') |
|---|---|---|
| *APP* | GCTGCCCAGCTTGGCACTGC | GGCAACGGTAAGGAATCACGATGTGGGTG |
| *PSD-95* | TCTGTGCGAGAGGTAGCAGA | AAGCACTCCGTGAACTCCTG |
| *Gadd45b* | GCTGGCCATAGACGAAGAAG | GCCTGATACCCTGACGATGT |
| *Zfyve1* | CTGGCAGTTCAAAAACTAGTAC | CCGGGGACAGCACCAATGAAAG |
| *Ap3d1* | CAAGGGCAGTATCGACCGC | GATCTCGTCAATGCACTGGGA |
| *Cpt1c* | AGAAGTAGAGCTCAGCTCGCCA | CCAGAGATGCCTTTTCCAGGAG |
| *Lamp2* | ATGTGCCTCTCTCCGGTTAAA | GCAAGTACCCTTTGAATCTGTCA |
| *Foxo6* | AAGAGCTCCCGACGGAAC | GGGGTCTTGCCTGTCTTTC |
| *TNF-α* | CTCATGCACCACCATCAAGG | ACCTGACCACTCTCCCTTTG |
| *IL-10* | CCCTTTGCTATGGTGTCCTT | TGGTTTCTCTTCCCAAGACC |
| *16S rDNA* | AGAGTTTGATCCTGGCTCAG | CTGCTGCCTCCCGTAGGAGT |
| *Gapdh* | TGGAGAAACCTGCCAAGTATGA | TGGAAGAATGGGAGTTGCTGT |
